# Modular *in vitro* evaluation of Buparlisib-polymeric nanomedicines in 2D and 3D models of glioblastoma

**DOI:** 10.64898/2026.06.18.732941

**Authors:** Jarmila Havelkova, Yuriy Petrenko, Anna Stehlíkova, Dana Marekova, Katerina Peskova, Michal Pechar, Martin Studenovsky, Tomas Etrych, Robert Pola, Pavla Jendelova

## Abstract

**Introduction:** In this study, we developed a modular in vitro platform that integrates advanced polymer–drug conjugation chemistry with stepwise cytotoxicity screening in both 2D (monolayer) and 3D (spheroids) glioblastoma (GBM) models. Buparlisib was selected as the model therapeutic due to its well-characterised mechanism of action, high blood–brain barrier permeability, and relevance to PI3K-targeted therapy.

**Methods:** Two mechanistically distinct conjugation strategies were explored using N-(2-hydroxypropyl)methacrylamide-based copolymers. The first strategy was based on a redox-sensitive disulphide linkage designed for intracellular glutathione-triggered release, whereas the second used an azide-bearing derivative compatible with strain-promoted azide–alkyne cycloaddition. Drug release was assessed by high-performance liquid chromatography. Biological activity was systematically evaluated in U87MG, U118MG, and T98G cells under 2D conditions using a resazurin-based metabolic activity assay. Subsequently, the more promising disulphide-based formulations were assessed in 3D spheroids by metabolic activity measurements and live-cell monitoring of spheroid growth dynamics.

**Results:** Free Buparlisib showed the strongest inhibitory effect, while its modification and polymer conjugation reduced the apparent activity. Nevertheless, the disulphide-based derivative and polymer conjugate retained concentration-dependent activity, whereas the azide-based polymer conjugate showed minimal effects. Moreover, treatment responses differed between cell lines and between 2D and 3D models.

**Discussion:** Overall, linker chemistry, cell-line-specific behaviour, and model dimensionality strongly influenced the biological performance of the polymeric Buparlisib formulations. The redox-sensitive polymer conjugate therefore represents the more promising strategy for further development.

## 1 Introduction

Glioblastoma (GBM) is the most aggressive and lethal primary brain tumour in adults, characterised by rapid progression, extensive heterogeneity, and resistance to conventional therapies (Louis et al., 2016). Standard-of-care treatment, including surgical resection, radiotherapy and temozolomide (TMZ) chemotherapy, offers only modest survival benefits, and recurrence is nearly universal (Tan et al., 2020). A major limitation of current therapeutic strategies is the poor penetration of drugs across the blood–brain barrier (BBB) and the inability to selectively target aberrant signalling pathways that drive GBM proliferation, invasion, and resistance (Noorani and de la Rosa, 2023, Wen et al., 2019). Among these, the phosphatidylinositol 3-kinase (PI3K)/Akt/mTOR pathway is frequently dysregulated in GBM, contributing to tumour growth, survival, and treatment resistance (Glaviano et al., 2023, Pournajaf and Pourgholami, 2025). Buparlisib (Bup), an orally available pan-PI3K inhibitor, has demonstrated potent anticancer activity in preclinical studies and excellent BBB permeability, yet its clinical application has been hampered by systemic toxicity, a narrow therapeutic window, and dose-limiting side effects (Netland et al., 2016, Speranza et al., 2016).

To overcome these challenges, polymer-based drug delivery systems have emerged as promising platforms for improving therapeutic index through enhanced solubility, tumour-selective accumulation, and controlled intracellular drug release (Tavares et al., 2021, Chytil et al., 2018, Parashar et al., 2024, Duncan, 2006). Importantly, the attachment of the drug to the delivery system often results in its complete inactivation, thereby minimising off-target side effects. After reaching the target zone, the following stimuli-sensitive degradation of the linker elicits a release of the drug from the delivery system, providing the desired effect. However, the efficiency of such systems strongly depends on the type of polymer carrier and the design of the linkage between the drug and carrier used for drug conjugation and release of the drug. In the case of Bup, which possesses multiple functional groups suitable for developing the derivatives, the selection of optimal linker chemistry is crucial to preserve activity, ensure intracellular activation, and avoid premature inactivation (Wang et al., 2025). However, despite the extensive preclinical and clinical evaluation of Bup, its chemical modification and incorporation into polymeric drug conjugates remain largely unexplored. In particular, no systematic comparison of different Bup linker and conjugation strategies within a single polymeric carrier has been reported to date. Moreover, the extent to which different conjugation strategies preserve Bup biological activity while enabling effective intracellular release remains unexplored and requires systematic experimental validation.

Equally critical to the predictive value of *in vitro* screening is the choice of an appropriate experimental model. Traditional two-dimensional (2D) cultures often fail to recapitulate the complexity of tumour architecture, extracellular matrix interactions, and drug diffusion barriers observed *in vivo*. In contrast, three-dimensional (3D) spheroid models more closely mimic the physiologic conditions of solid tumours, including gradients of oxygen, nutrients, and therapeutic agents, as well as multicellular resistance phenotypes (Nayak et al., 2023, Abbas et al., 2023). Nevertheless, the compatibility of different GBM cell lines with 3D culture formats and their differential responses to drug formulations can vary significantly depending on their genetic background and intrinsic metabolic states.

This study provides the first systematic comparison of two chemically distinct strategies for Bup modification and conjugation within a single polymeric carrier, together with direct evaluation of the resulting constructs in both 2D and 3D *in vitro* models (classic monolayer and spheroid cultures, respectively). We developed a modular *in vitro* platform that integrates advanced polymer–drug conjugation chemistry with stepwise cytotoxicity screening in both 2D and 3D GBM models. Bup was selected as the model therapeutic due to its well-characterised mechanism of action, high BBB penetration, and relevance to PI3K-targeted therapy. Two mechanistically distinct conjugation strategies were explored for the synthesis of a drug delivery system based on the *N*-(2-hydroxypropyl)methacrylamide-based copolymers (pHPMA). The first strategy is based on a redox-sensitive disulphide linkage designed for intracellular glutathione-triggered release. The second approach is based on an azide-bearing derivative compatible with strain-promoted azide–alkyne cycloaddition (SPAAC) for biorthogonal click conjugation to polymeric carriers. Although both approaches enabled stable polymer attachment and potential intracellular drug liberation, their biological efficiency, release kinetics, and compatibility with the target cell environment were unknown and required systematic evaluation. Their biological activity was then systematically evaluated in three GBM cell lines (U87MG, U118MG, T98G), mimicking the differential treatment response. The use of multiple model systems enabled a comparative assessment of drug efficacy, linker performance, and cell-line-specific behaviour in both 2D and 3D settings.

## 2 Materials and methods

### 2.1 Chemicals

Methacryloyl chloride, 3-aminopropanoic acid (β-Ala), 4,5-dihydrothiazole-2-thiol, dimethylaminopyridine, 1-ethyl-3-(3-dimethylaminopropyl)carbodiimide hydrochloride, azobisisobutyronitrile (AIBN), 4-cyano-4-thiobenzoylsulfanylpentanoic acid, tert-butyl alcohol (t-BuOH), *N,N*-dimethylacetamide (DMA), tris(2-carboxyethyl)phosphine (TCEP), glutathione (GSH) were purchased from Sigma-Aldrich. *N,N*-diisopropylethylamine (DIPEA) and 2-(*N*-Boc-2-aminoethyldisulfanyl)ethanol were purchased from Iris Biotech, Germany. Dibenzocyclooctyne-amine (DBCO-NH_2_) and 3-azidopropanol were purchased from Click Chemistry Tools, USA. Acetonitrile, dichloromethane, methanol and other common solvents and chemicals were purchased from Merck, s.r.o. Bup was purchased from MedChemExpress, Sollentuna, Sweden. 1-Amino-2-propanol (AMP) was purchased from TCI, Tokyo, Japan. Initiator V-70 was purchased from FUJIFILM Wako Chemicals, USA. All chemicals and solvents were of analytical grade.

### 2.2 Cell culture reagents and materials

MEMα media with GlutaMAX™ supplement (Gibco™, United Kingdom, cat. no. 32561029), Dulbecco’s Modified Eagle Medium/Nutrient Mixture F-12 (DMEM/F-12) medium (Gibco™, United Kingdom, cat. no. 31331028); Fetal Bovine Serum (FBS) (Collected in South America, Capricorn Scientific, Germany, cat. no. FBS-12A); Penicillin-Streptomycin (100X) (Gibco™, USA, cat. no. 15070–063), Resazurin sodium salt (Sigma®, cat. no. 62758-13-8).

Cultivation flasks (Nunc™ EasYFlask™ Cell Culture Flask; Thermo Scientific™, cat. no. 156499), flat-bottom 96-well plate (Tissue Culture Testplate 96F; TPP™, cat. no. 92096), BIOFLOAT™ 96-well plate (faCellitate GmbH, Germany, cat. No. F202003), Incucyte® Sx5 Live-Cell Analysis Instrument (Incucyte® Sx5, Sartorius, Germany) -compatible ultra-low attachment U-shape 96-well plate (Costar®, Corning®, USA, cat. no. 7007)

### 2.3 Chemical analysis

An HPLC equipped with reversed-phase Chromolith Performance RP-18e columns (100×4.6 mm, Merck, Germany) and with a UV-VIS diode array detector (Shimadzu, Japan) was used to monitor the synthesis and purity of the monomers. The mobile phase consisted of a linear gradient of water/acetonitrile (0-100 % acetonitrile) in the presence of 0.1 % TFA. The molecular mass of the derivatives was determined using mass spectrometry performed on an LCQ Fleet mass analyser with electrospray ionisation (ESI-MS) (Thermo Fisher Scientific, Inc., MA, USA). Molecular weights and dispersity of the copolymers and the drug release from the polymer conjugate were determined by size exclusion chromatography (SEC) on a HPLC system (Shimadzu, Japan) equipped with the following detectors: UV, differential refractive index, multi-angle light scattering (LS) DAWN Helleos II and viscosimetric detector ViscoStar III (all Wyatt Technology Corp., USA). The analysis was performed using either a TSKgel G3000SWXL column (Tosoh Bioscience, Japan) (methanol/phosphate buffer, 8/2, pH 6.5) at a flow rate of 0.5 mL·min^−1^ or Superose 6 Increase 10/300 GL (Cytiva, USA) (0.2 M phosphate buffer) at a flow rate of 0.5 mL·min^−1^. The contents of the thiazolidine-2-thione (TT) groups DBCO groups were determined on a Helios Alpha UV/VIS spectrophotometer (Thermospectronic, UK) using the absorption coefficients for TT in methanol (ε_305_=10,300 L mol^−1^ cm^−1^) and for DBCO also in methanol (ε_292_=13,000 L·mol^−1^·cm^−1^).

### 2.4 Synthesis of 3-azidopropyl carbamate derivative of Buparlisib (AP-Bup)

3-Azidopropanol (100 µL, 1.09 mmol) was mixed with phosgene (3 mL of 20% solution in toluene, 5.7 mmol) and kept under an argon atmosphere at 4 °C for 4 days (for the scheme of synthesis see **Supplementary Fig. 1**). The excess of phosgene and solvent were subsequently removed by a gentle stream of argon under heating at 30 °C, followed by brief evacuation under reduced pressure. The resulting 3-azidopropyl chloroformate was obtained as a colourless liquid and used in the next step without further purification. Then, a solution of Bup (22 mg, 53.6 µmol) in dichloromethane (1 mL) and DIPEA (11.2 µL, 64.3 µmol) were added, and the reaction mixture was stirred at room temperature. According to HPLC monitoring, the conversion was complete within 20 min. The mixture was diluted with hexane and concentrated under reduced pressure. The resulting precipitate was repeatedly washed with hexane and dried under vacuum. The crude product was dissolved in acetone, filtered through a PTFE syringe filter, and crystallised by slow evaporation from hexane to afford a white solid. The crude product was dissolved in dichloromethane (20 mL) and extracted three times with 30 mL of water. The combined organic layers were dried over anhydrous sodium sulfate, filtered, and concentrated under reduced pressure. Crystallisation was induced by the addition of an excess of hexane, followed by partial solvent evaporation. The obtained white solid was collected by filtration and dried in a vacuum. Yield: 19 mg (66%). ESI-MS: calculated 537.2, found 538.1 [M+H]⁺ (for the spectrum see **Supplementary Fig. 2**). ^1^H NMR (400 MHz, CDCl_3_): 8.52-8.43 (br s, 1 H, Ar); 8.41-8.34 (br s, 1 H, Ar); 6.07-5.98 (br s, 1 H, Ar); 4.39-4.30 (t, 2 H, *J* = 6.3 Hz, -C*H_2_*-O-C(=O)-); 3.98-3.56 (m, 16 H, -N-C*H_2_*-C*H_2_*-O-); 3.49-3.43 (t, 2 H, *J* = 6.6 Hz, -N_3_-C*H_2_*-CH_2_-); 2.05-1.96 (p, 2 H, *J* = 6.3 Hz, -N_3_-CH_2_-C*H_2_*-CH_2_-O-) ppm.

### 2.5 Synthesis of 2-(2-aminoethyldisulfanyl)ethyl derivative of Buparlisib (SS-Bup)

2-(*N*-Boc-2-aminoethyldisulfanyl)ethanol (117 mg, 0.198 mmol) was dissolved in phosgene (3 mL of 20% solution in toluene 5.7 mmol), DIPEA was added (163 µL, 0.94 mmol) and the reaction mixture was stirred 30 min at RT. The excess phosgene and solvent were subsequently removed by a gentle stream of argon under heating at 30 °C, followed by brief evacuation under reduced pressure. Then, a solution of Bup (66 mg, 0.16 mmol) in dichloromethane (3 mL) and DIPEA (150 µL; 0.86 mmol) were added, and the reaction mixture was stirred at room temperature for 30 min. The solvents were removed under reduced pressure, and TFA (2 mL) was added to remove the protecting groups from the amino group (for the scheme of synthesis see **Supplementary Fig. 3**). The deprotection proceeded instantaneously. The crude product was purified by preparative HPLC on a C18 reversed-phase column using a linear gradient of 0–100% acetonitrile/water containing 0.1% TFA. The collected fractions were combined, and the solvents were removed under reduced pressure. The purified product was obtained as a solid and dried under vacuum. Yield: 59 mg (62%). ESI-MS: calculated 589.2, found 590.1 [M+H]⁺. ^1^H NMR (400 MHz, DMSO-*d6*): 10.93(s, 1 H, -N*H*-); 8.55 (s, 1 H, Ar); 8.26 (s, 1 H, Ar); 8.03-7.72 (br s, 3H, -N*H_3_*^+^); 6.37 (s, 1 H, Ar); 4.45-4.38 (t, 2 H, *J* = 6.5 Hz, -NH-C(=O)-O-C*H_2_*); 3.71-3.55 (m, 16 H, -N-C*H_2_*-C*H_2_*-O-); 3.18-3.04 (m, 4 H, -C*H_2_*-S-S-C*H_2_*-); 2.99-2.92 (m, 2 H, -C*H_2_*-NH_3_^+^) ppm.

### 2.6 Synthesis of the monomers

*N*-(2-Hydroxypropyl)methacrylamide (HPMA) was prepared as described previously (Ulbrich et al., 2000). Methacrylamidopropanoic acid (Ma-β-Ala-OH) was prepared by the Schotten-Baumann acylation procedure as previously described^1^. 3-(3-Methacrylamidopropanoyl)thiazolidine-2-thione (Ma-β-Ala-TT) was prepared as described previously (Pola et al., 2021). The monomer was characterised using HPLC (single peak), ^1^H NMR spectra (**Supplementary Fig. 4**) and ESI-MS (Ma-β-Ala-TT: calculated 258.3, found 281.1 [M+Na]).

### 2.7 Synthesis of the polymer precursors

Polymer precursor **Prec1**, poly(HPMA-*co*-Ma-□-Ala-TT), was prepared by reversible addition-fragmentation chain transfer (RAFT) copolymerisation of HPMA (88 mol%, 100 mg) and Ma-□-Ala-TT (12 mol%; 24.6 mg) using 4-cyano-4-thiobenzoylsulphanylpentanoic acid (0.54 mg) as a chain transfer agent (CTA) and V-70 (0.38 mg) as a low temperature initiator. The molar ratio of monomers:CTA:initiator was 650:2:1. The polymerisation was carried out in the mixture of t-BuOH/DMA – 85/15 v/v (1.1 mL; 0.7 M solution of monomers) and transferred into a glass ampule where the mixture was bubbled with Ar and sealed. After 16 h at 40 °C, the product was precipitated into diethyl ether:acetone (1:1); the precipitate was then washed with diethyl ether and dried under vacuum. Polymer precursor was then reacted with AIBN (10 molar excess) in DMA (15 % w/w solution of polymer) under Ar for 2 h at 70 °C in a sealed ampule to remove the dithiobenzoate (DTB) ω-end groups (Perrier et al., 2004). The reaction mixture was precipitated into diethyl ether:acetone (1:1). The precipitate was washed with diethyl ether and dried under vacuum to yield polymer precursor **Prec1**.

Polymer precursor **Prec2** was prepared by the reaction of DBCO-NH_2_ (11.8 mg, 42.7 □mol) with **Prec1** (168.25 mg) in the ratio 11.7 to 4 (TT to DBCO groups) in the presence of 11 □L DIPEA (64.3 □mol). The course of reaction was observed by HPLC, and after 2 hours, the reaction was ended by the addition of AMP (10 µL). The solution of polymer was precipitated in acetone/diethyl ether (1:1), washed with diethyl ether and dried to yield 159 mg (88 %) of precursor **Prec2** as a white powder.

The characterisation of the prepared polymer precursors is shown in **Table 1** and NMR spectrum and SEC chromatogram of polymer precursor **Prec1** see **Supplementary Fig. 4 and 5A.**

**Table 1.**
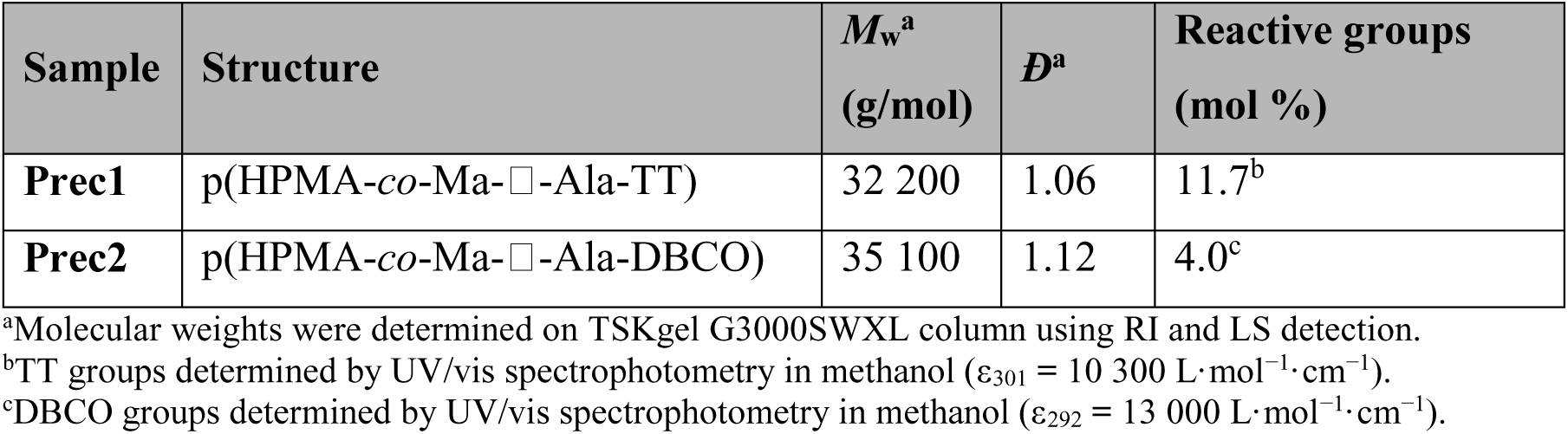
Characterisation of the polymer precursors.

### 2.8 Synthesis of polymer conjugate P-SS-Bup

Conjugate **P-SS-Bup** was prepared by conjugation of 9.75 mg of Bup-SS-NH₂*TFA (13.9 µmol), corresponding to 17.2 wt% of the total derivative or 10 wt% of free **Bup** (without the spacer SS-NH_2_*TFA), with the polymer precursor **Prec1** (46.9 mg) in MeOH (0.5 mL) in the presence of 4.8 □L DIPEA (41.7 □mol).

The course of the reaction was observed by HPLC. After 4 hours when no free Bup-SS-NH_2_ was detected, the reaction was ended by the addition of AMP (2.8 µL) (for synthetic scheme and HPLC chromatograms of the reaction see **Supplementary Fig. 6**). The crude product was precipitated into acetone/diethyl ether (1:1), and the precipitate was washed with diethyl ether and dried. The polymer was dissolved in water, purified by chromatography on Sephadex G 25 resin in water (PD 10 column, Pharmacia), and freeze-dried to yield 46.5 mg (85 %) of conjugate **P-SS-Bup** as a white powder. The characterisation of the prepared polymer precursors is shown in **Table 2** and SEC chromatogram of polymer conjugate **P-SS-Bup** see **Supplementary Fig. 5B.**

**Table 2.** Characterisation of the polymer conjugates on Superose 6 Increase 10/300 GL.

| Sample | Structure | $M_w^a$<br>(g/mol) | $\bar{D}^a$ | $D_h^a$<br>(nm) |
| --- | --- | --- | --- | --- |
| **P-SS-Bup** | p(HPMA-co-Ma-□-Ala-NH-SS-Bup) | 54 700 | 1.41 | 12.1 |
| **P-AP-Bup** | p(HPMA-co-Ma-□-Ala-NH-DBCO-AP-Bup) | 42 600 | 1.25 | 12.4 |

<sup>a</sup>Molecular weights and hydrodynamic diameter were determined by SEC using RI, LS and viscosimetric detection.

### 2.9 Synthesis of polymer conjugate P-AP-Bup

Conjugate **P-AP-Bup** was prepared by conjugation of 7.7 mg of AP-Bup (14.3 mol, 13.1 wt% corresponding to 10 wt% of free of Bup) with reactive polymer precursors **Prec2** (51 mg) in DMA (0.5 mL) (for synthetic scheme see **Supplementary Fig. 7**). The course of the reaction was observed by HPLC, and after 1.5 hours, there was no free **AP-Bup**. The reaction mixture was added dropwise into acetone/diethyl ether (1:1), and the precipitate was washed with diethyl ether and dried. The polymer was dissolved in water, purified by chromatography on Sephadex G 25 resin in water (PD 10 column, Pharmacia), and freeze-dried to yield 52.4 mg (89 %) of conjugate **P-AP-Bup** as a white powder. The characterisation of the prepared polymer precursors is shown in **Table 2**.

### 2.10 Release of Bup from P-SS-Bup or P-AP-Bup from polymer conjugate at pH 7.4

Polymer conjugate **P-SS-Bup** (equivalent to 100 µM or 50 µM of Bupwas dissolved in 75 mM tris-HCl buffer (pH 7.4). The release was initiated by adding a GSH stock solution (100 mM) to obtain a final GSH concentration of 1 mM. The total reaction volume was 450 µL, and the samples were incubated at 25 °C. At predetermined time intervals, 10 µL aliquots were withdrawn, and the reaction was quenched by adding 5 µL of 6% aqueous trifluoroacetic acid (TFA). The amount of Bup released was quantified by HPLC. The current evaluation system represents a simplified chemical model for verifying the functionality of the redox-sensitive disulphide linker and confirming the formation of native Bup under reducing conditions. Therefore, the presented release kinetics should be considered an estimate, as drug release is expected to proceed more rapidly under physiological temperature and intracellular conditions.

The spontaneous release of Bup from **P-SS-Bup** or **P-AP-Bup** in buffer with pH 7.4. was performed as follows: Polymer conjugate (equivalent to 100 µM or 50 µM of Bup) was dissolved in 75 mM tris-HCl buffer (pH 7.4). The total reaction volume was 450 µL. At 12 h and 24 h intervals, 10 µL aliquots were withdrawn, and the reaction was quenched by adding 5 µL of 6% aqueous trifluoroacetic acid (TFA). The amount of Bup released was quantified by HPLC.

For complete (100%) release, the same procedure was performed using tris-(2-carboxyethyl)phosphine (TCEP) (**Supplementary Fig. 8)** instead of GSH at an equimolar concentration. The reaction progress was monitored by HPLC, and the area of Bupb peak at the point when no intermediate cleavage products were detected was taken as 100% release. In the early stages, additional peaks corresponding to intermediate species were observed, which gradually disappeared as a single Bup peak became predominant. This procedure was adapted from the methodology previously described (Neumann and Nolan, 2018).

### 2.11 Cell culture and cell seeding

The human glioblastoma cell lines used in this study were U87MG (ATCC® HTB-14™, LGC Standards Sp. Z.o.o., Poland), U118MG (ATCC® HTB-15™, LGC Standards Sp. Z.o.o., Poland) and T98G (ATCC® CRL-1690™, LGC Standards Sp. Z.o.o., Poland). Upon receipt, cells underwent a quarantine period in a separate incubator and were tested for mycoplasma before being incorporated into any experimental workflow.

All cell cultures were tested for mycoplasma contamination using Venor®GeM qEP (Minerva Biolabs®) real-time PCR (qPCR) assay, according to manufacturer’s instructions, and were confirmed to be mycoplasma-free prior to use in experiments. We acknowledge the potential risk of cross-contamination from externally sourced cell lines and took appropriate precautions.

All GBM cell lines were cultured in cultivation flasks in complete cell culture medium (CCM) (DMEM/F-12 medium supplemented with 10 % FBS, and 1X Penicillin-Streptomycin (final concentration in CCM: 50 units/mL Penicillin, 50 µg/mL Streptomycin) in a humidified 5 % CO_2_ atmosphere at 37 °C.

Due to the different proliferation rates of each cell line, for 2D experiments, cells were seeded at 70% confluency in a flat-bottom 96-well plate to prevent overgrowth during the treatment period (for details see Treatment regimen section). Seeding densities for cell lines U87MG, U118MG, and T98G were 12 × 10^3^, 14 × 10^3^, and 10 × 10^3^ per well (growth area of 0.345 cm^2^), respectively. Following seeding, cells were cultured under standard conditions overnight to allow for proper attachment before treatment.

For 3D experiments, cells (20 × 10^3^/well) were seeded in the BIOFLOAT™ 96-well plate (200 µL CCM/well) and cultured under standard conditions for 3 days to allow for formation of compact, circular spheroids. At day 3 of cultivation, spheroids were carefully transferred to an Incucyte® Sx5-compatible imaging vessel for treatment and subsequent response monitoring.

### 2.12 Treatment regimen

#### 2.12.1 2D cell cultures

One day prior to treatment, cells were seeded as described in the previous section (2.11. Cell culture and cell seeding) and incubated overnight to allow for proper attachment. Following day (D0; see **Supplementary Fig. 9A**), cells were treated with selected substances as follows: free Bup, **SS-Bup**, and **P-SS-Bup** at several concentrations (2, 5, 20, 50, 100, and 200 µM) (**Fig. 1**), or free Bup, **AP-Bup**, and **P-AP-Bup** at concentrations 50 µM and 200 µM (**Fig. 3**). Each biological replicate was performed with a minimum of two technical replicates. The treatment duration was 72 hours (see **Supplementary Fig. 9A**). The concentrations (µM) refer to the actual concentration of Bup in the well. For the negative control groups, CCM was used without DMSO and with DMSO at concentrations corresponding to those used to dissolve the tested substances to final concentrations of 50 µM and 200 µM. After 3 days (D3), an adjusted resazurin assay was performed to measure the metabolic activity of all cells (attached, loose) in wells (see section 2.14.1 Metabolic activity measurement – 2D cell cultures).

**Figure 1.**
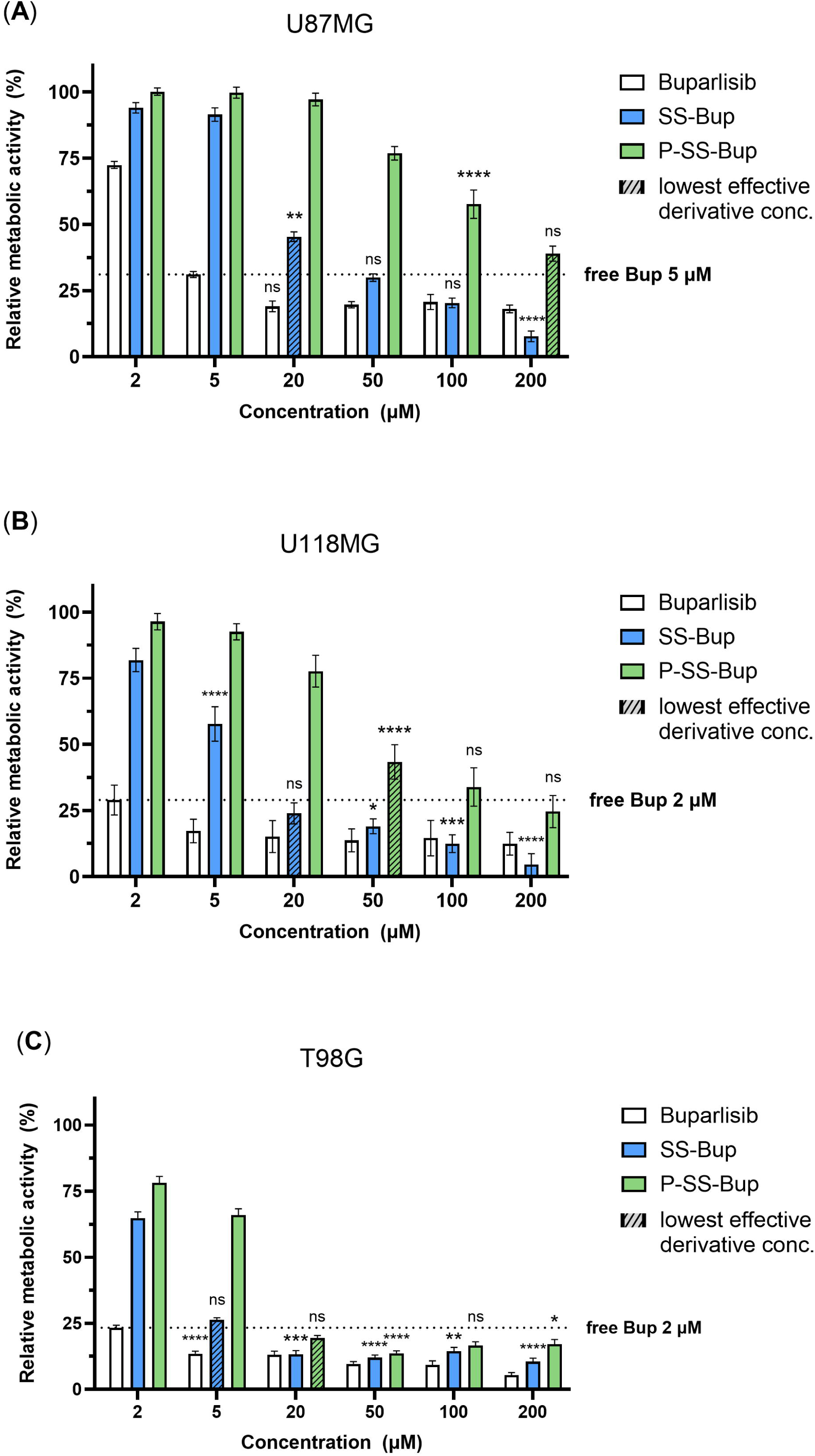
Effect of drug modification (SS) on the metabolic activity of GBM cell lines in 2D conditions. Comparison of metabolic activity of 2D-cultured U87MG (**A**), U118MG (**B**), and T98G (**C**) glioblastoma cell lines after a 72-hour treatment with free Bup (white), pro-drug containing a disulphide bridge (**SS-Bup**; blue), and polymer-conjugated drug delivery construct **(P-SS-Bup**; green). Values were normalised to the untreated control group (set to 100 %). Dotted horizontal lines indicate the mean values of the lowest effective free Bup concentration, which were used for the statistical comparison of all groups. Data presented as mean ± SEM. Significance levels (adjusted p - values): [ns] (p_adj ≥ 0.05); [*] (p_adj < 0.05); [**] (p_adj < 0.01); [***] (p_adj < 0.001); [****] (p_adj < 0.0001). n=4-9

#### 2.12.2 3D cell cultures

Spheroids were prepared as described in the section 2.11. Cell culture and cell seeding. Upon transfer to an Incucyte® Sx5**-**compatible imaging vessel at day 3 of cultivation (D0 of treatment period; see **Supplementary Fig. 9B**), formed spheroids were treated with selected substances (free Bup, **SS-Bup**, **P-SS-Bup)** at selected concentrations (50 and 200 µM). Each biological replicate was performed with a minimum of three technical replicates. For the negative control groups, CCM was used without DMSO and with DMSO at concentrations corresponding to those used to dissolve the tested substances to final concentrations of 50 µM and 200 µM. Next, the plate was placed into Incucyte® Sx5 Live-Cell Analysis Instrument (Incucyte® Sx5; for details see the next section) and monitored for the duration of 60–62 hours (D3, see **Supplementary Fig. 9B**). After the treatment period, an adjusted resazurin assay (see the section 2.14.2 Metabolic activity measurement – 3D cell cultures) was performed to measure the metabolic activity of spheroids in response to the treatment.

### 2.13 Spheroid growth dynamics monitoring

For monitoring the spheroid growth dynamics, the Incucyte® Sx5 instrument was utilised. Upon placement of the imaging vessel, wells were scanned every 2 hours (image acquisition). Image segmentation and analysis were performed using Incucyte® Sx5 built-in software, and subsequent analysis of spheroid growth dynamics was performed in Excel/GraphPad Prism. To investigate differences between cell lines, untreated spheroid monitoring cultivation time in the Incucyte® Sx5 was set to 46 hours.

### 2.14 Metabolic activity measurement

Metabolic activity of cells was monitored using the resazurin assay. Resazurin stock solution was prepared as follows: 5.5 mg of resazurin powder (Resazurin sodium salt) was dissolved in 10 ml of MEMα media, filtered, and diluted to a final volume of 50 ml. For performing the assay, the stock was diluted to a final concentration of 10 % in complete cell culture medium.

The resazurin assay was adjusted for 2D and 3D experiments as follows.

#### 2.14.1 2D cell cultures

After the treatment period (D3, see **Supplementary Fig. 9A**), part of the medium in the wells was discarded to prevent cell detachment or loss, and resazurin solution was added at a final concentration of 10 %. Wells designated for background correction contained CCM with resazurin solution at a corresponding concentration (10 %). After 2 hours of incubation in a humidified 5 % CO_2_ atmosphere at 37 °C, 200 µL of resazurin solution from each well was transferred to a new flat-bottom 96-well plate for fluorescence measurement in a TECAN GENios microplate reader (Tecan) with the following settings: excitation wavelength 550 nm, emission wavelength 590 nm.

#### 2.14.2 3D cell cultures

After the treatment period (D3, see **Supplementary Fig. 9B**), part of the medium in the wells was discarded, and resazurin solution was added (10 % final concentration). Wells designated for background correction contained CCM with resazurin solution at a corresponding concentration (10 %). After 4 hours of incubation in a humidified 5 % CO_2_ atmosphere at 37 °C, 100 µL of resazurin solution from each well was transferred to a new flat-bottom 96-well plate for fluorescence measurement in a TECAN microplate reader (settings: excitation wavelength 550 nm, emission wavelength 590 nm).

### 2.15 Spheroid immunocytofluorescence staining

To inspect the viability of cells in 3D cultures, spheroids were washed with PBS and incubated in HBSS solution with live/dead cell viability dyes propidium iodide (PI) and fluorescein diacetate (FDA) for 10-15 minutes. Before image acquisition, spheroids were washed twice with PBS. Image acquisition was performed using a versatile spinning disc confocal microscope Nikon CSU W1 with a 10X objective.

### 2.16 Statistical analysis

In all experiments, the value “*n*” refers to number of independent biological replicates of selected groups, each performed with a minimum of two technical replicates. If a range of “*n*” is depicted, e.g. n=4–9, the lower number refers to the minimum number of biological replicas of selected groups, and similarly, the higher number the maximal number of biological replicas of selected groups.

As DMSO, used as a solvent for tested substances, did not cause a significant change in metabolic activity in 2D conditions or in spheroid growth dynamics in 3D conditions (see Supplementary Fig. 10A and 10B and C, respectively), for all comparative experiments, the values for control group were averaged from untreated sample (CCM only), and samples treated with solvent (DMSO) at concentration corresponding to the solvent concentration in groups with 50 µM and 200 µM derivatives.

#### 2.16.1 Resazurin assays

To identify the lowest effective concentrations of Bup and other substances, for each experimental dataset, raw data from the resazurin assays were compared to the averaged values from control groups using Welsch’s ANOVA method, followed by Dunnett’s T3 post-hoc test in GraphPad Prism, with p-value adjustment (**Supplementary Table 1a, 1b**). For clarity of further statistical comparisons between groups, results of this analysis are only commented on in the Results section, and graphically depicted in **Figure 1** (A, B, C) as either a dotted line (the lowest effective concentration of Bup) or a hatched bar (the lowest effective concentration of Bup derivatives).

Prior to a comparison of the effectiveness of Bup derivatives to the effective concentrations of free Bup, the assumptions of the ANOVA model were inspected. Subsequently, datasets were analysed using either the Aligned Rank Transform (ART) ANOVA in the ARTool package (v0.11.2; (Wobbrock et al., 2011, Kay et al., 2025) in R (v4.3.0) (assumptions violated; 2D metabolic activity) or the One-way ANOVA method in GraphPad Prism (assumptions met; 3D metabolic activity). For comparative statistical tests performed on normalized data from 2D resazurin assays, corrected values were normalised to the averaged control groups (defined as 100 %) and are presented as mean ± SEM. For 3D resazurin assays, data are presented as mean Relative Fluorescent Units (RFU) ± SEM.

For multiple comparisons of tested groups following the One-way ANOVA method, the post-hoc analysis was performed using Tukey’s Honestly Significant Difference (HSD) test, and Tukey’s p-value adjustment was applied to correct for multiple comparisons. Differences were considered statistically significant at *p* < 0.05.

The ART method was selected because the data did not meet the assumptions of normality and homogeneity of variance required for parametric ANOVA (Wobbrock et al., 2011), and standard non-parametric tests (e.g., Kruskal-Wallis) do not validly assess interaction effects in multifactorial designs. Two-way factorial design was implemented with Treatment and Concentration as fixed factors. Post-hoc pairwise comparisons for the interaction effect were conducted using the art.con function (Elkin et al., 2021). To maintain statistical power, contrasts were performed on the aligned responses with p-values adjusted for multiple comparisons using Tukey’s HSD method. Differences were considered statistically significant at *p* < 0.05.

Prior to all statistical tests, the measured values from the TECAN microplate reader were first background-corrected.

#### 2.16.2 Spheroid growth dynamics

Data from each individual spheroid obtained from Incucyte® Sx5 Live-Cell Analysis Instrument were normalised to its starting timepoint value (0h) and analysed using a mixed-effects model (REML; GraphPad Prism) to account for repeated measurements from the same spheroids over time, with time and treatment as fixed effects and spheroid biological replica ID as a random effect. Multiple comparisons between experimental groups at individual time points were performed using GraphPad Prism built-in procedures (Tukey’s test) to control for multiple testing. Data are presented as means (SEM omitted in some cases for visual clarity of the charts; see **Supplementary Material 3**), and differences were considered statistically significant at *p* < 0.05.

## 3 Results

To enable the incorporation of Bup into polymer–drug conjugates, two reactive derivatives were designed, differing in the type of cleavable linkage incorporated into the derivative structure. The rationale behind this design was to achieve controlled intracellular release of the active drug under distinct biological stimuli. The first derivative (**SS-Bup**) incorporates a disulphide bridge that is sensitive to reductive cleavage by intracellular GSH, allowing drug release specifically in the reductive environment of the cytosol. The second derivative (**AP-Bup**) contains a potentially enzymatically cleavable carbamate bond, which is expected to undergo hydrolysis by lysosomal enzymes after the endocytosis of the conjugate (Di, 2019, Ghosh and Brindisi, 2015). This dual design enables a direct comparison of two distinct biodegradation mechanisms and their influence on the release kinetics and biological activity of the polymer-Bup conjugates.

### 3.1 Synthesis of Buparlisib derivatives SS-Bup and AP-Bup

In both synthetic routes, suitable alcohols served as starting materials: 2-(*N*-Boc-2-aminoethyldisulfanyl)ethanol for **SS-Bup** and 3-azidopropanol for **AP-Bup**. In the first step, the alcohols were treated with an excess of phosgene in toluene and formed the corresponding chloroformates. These reactive intermediates were then successfully coupled with Bup in the presence of a base to yield the respective derivatives. The **AP-Bup** derivative was obtained directly, while the **SS-Bup** derivative was initially isolated in its Boc-protected form and subsequently deprotected with trifluoroacetic acid (TFA) to afford the desired compound. Both derivatives were obtained in moderate yields of approximately 60% and exhibited high purity as confirmed by analytical HPLC. The chemical structures of the compounds were verified by ¹H NMR spectroscopy and mass spectrometry, which confirmed the expected molecular ion peaks corresponding to the respective derivatives.

### 3.2 Synthesis of polymer conjugates with SS-Bup and the effect of drug modification on the metabolic activity of GBM cell lines in 2D conditions

Prior to the synthesis of the polymer-drug conjugate **P-SS-Bup** the polymer precursor **Prec1** was successfully synthesised via RAFT copolymerisation of HPMA and Ma-β-Ala-TT. The prepared polymer has well-defined composition and a narrow dispersity (Đ = 1.06), indicating good control over the polymerisation process. Subsequent radical-induced cleavage of the dithiobenzoate end group using AIBN resulted in the removal of the CTA, which is critical for further bioconjugation and biological applications to minimise potential cross-linking and cytotoxicity associated with residual thiocarbonylthio-moieties.

The polymer-drug conjugate **P-SS-Bup** was then synthesised via efficient coupling of the amine-functionalised pro-drug **Bup-SS** to the reactive thiazolidine-2-thione (TT) groups of the polymer precursor **Prec1**, and its progress was monitored by HPLC to ensure the controlled conjugation of all **Bup-SS** present in the reaction mixture (see **Supplementary Fig. 6B**). After subsequent purification by precipitation and gel filtration chromatography, the final conjugate was obtained in high yield (85%) as a white solid, confirming the efficient removal of released TT groups and organic solvents. The final product contained 10 wt% of Bup, corresponding to the 17.2 wt% of Bup-SS-NH₂·TFA used in the initial reaction feed. The incorporation of a disulphide-containing (SS) spacer provides a reductively cleavable linkage, enabling drug release in response to the intracellular reducing environment.

To evaluate the impact of Bup modification through the pro-drug synthesis (**SS-Bup**) and subsequent conjugation to the HPMA polymer (**P-SS-Bup**), we performed a comparative analysis of free Bup and its derivatives on 2D cultures of three GBM cell lines: U87MG, U118MG, and T98G. Cells were treated with various concentrations of Bup formulations (2, 5, 20, 50, 100, and 200 µM, calculated for free Bup) for 72 hours. After the treatment period, metabolic activity was assessed using the resazurin assay.

In all three cell lines, 72-hour treatment with free Bup led to a significant, concentration-dependent reduction in metabolic activity. However, at lower concentrations (2–5 µM), the degree of suppression varied between cell lines, with U87MG showing approximately two-times higher metabolic activity at 2 µM concentration (≈ 28% decrease), statistically indifferent from its control. In contrast, U118MG and T98G cells showed a significant reduction in metabolic activity (approximately 75%) at the same concentration (**Fig**. **1B****, 1C**, respectively; **Supplementary Table 1a**). Increasing the concentration to 5 µM caused a significant reduction in metabolic activity of approximately 70% in U87MG cells (**Fig. 1A**), and an even more pronounced effect in the other two cell lines (**Fig. 1B, 1C**). Further increases in concentration did not significantly enhance the inhibitory effect in U118MG cells, indicating a maximum suppression (**Fig. 1B**). In T98G cells, only the highest concentration of 200 µM showed a significantly stronger effect than the 5 µM concentration (p_adj=0.0008) (**Fig. 1C**). Maximum suppression in U87MG cells was achieved at 20 µM (**Fig. 1A**; significance between free Bup concentration groups not shown for clarity of the following results).

Modification of the free Bup by introducing a disulphide (SS) linker led to a marked reduction in its ability to inhibit the metabolic activity of GBM cells. After 72 hours of treatment with 2 µM **SS-Bup**, we observed no significant effect on any of the cell lines, with minimum decrease of metabolic activity in U87MG cells (≈ 6%). Around 18% and 35% reductions of metabolic activity were observed in the case of the U118MG and T98G cells, respectively, compared to approximately 75% inhibition with free Bup at the same concentration (**Fig. 1B, 1C**).

Further analysis revealed that increasing **SS-Bup** concentrations resulted in a gradual, dose-dependent inhibition of metabolic activity across all three cell lines, however, the inhibition profiles varied. In the most sensitive cell line, T98G, a substantial inhibitory effect was observed at the second lowest concentration (5 µM), while reaching the maximal inhibitory effect at 20 µM concentration (≈87% decrease), which was statistically comparable to the 5 µM free Bup inhibition (≈86% decrease) (**Fig. 1C**). Further increases to 50 µM and beyond resulted in approximately 90% inhibition, with no additional benefit observed at higher concentrations. In U118MG cells, the inhibitory effect of **SS-Bup** developed slightly slower. While the 5 µM free Bup resulted in an 83% decrease in metabolic activity, the **SS-Bup** required a 20µM concentration to reach a comparable significant effect (≈76% decrease) (**Fig. 1B**). A similar trend was observed in U87MG cells, where exposure to **SS-Bup** resulted in even more concentration-wise delayed inhibition. In U87MG cells, the lowest concentration of **SS-**Bup resulting in a significant effect (≈55% decrease, p_adj=0.0088, significance not depicted in **Fig. 1A**, see **Supplementary Table 1a**) was 20 µM, while at 50 µM, the resulting inhibitory effect statistically matched the effect of 5 µM free Bup (≈69% decrease, p_adj=1) (**Fig. 1A**).

Therefore, the efficacy of the **SS-Bup** derivative was consistently lower than that of free Bup across all studied cell lines, with comparable effects observed only at concentrations approximately 4- to 10-fold higher (i.e., 5–20 µM **SS-Bup** vs. 2–5 µM free Bup; **Supplementary Table 1a**). These results demonstrate a dose-dependent effect of both substances, where increasing concentrations lead to a greater inhibition of metabolic activity in U118MG and U87MG cells. Although the molecular weight of **SS-Bup** is only about 50 % higher than free Bup, keeping it within the threshold for passive cellular diffusion, the pro-drug requires intracellular reduction by GSH to release the active Bup molecule. This activation step likely limits the immediate bioavailability of the drug inside cells, contributing to the attenuated inhibitory effects.

Similar trends were observed with **P-SS-Bup** treatment, although higher concentrations of the polymeric conjugate were required to achieve an inhibition comparable to that of free Bup in significantly effective concentration (**Fig. 1**; mind dotted lines). In U87MG cells, a concentration of 200 µM **P-SS-Bup** was necessary to match the inhibitory effect of 5 µM free Bup. In contrast, the response was stronger in U118MG and even more pronounced in T98G cells: 100 µM **P-SS-Bup** was sufficient to reach approximate the effect of 2 µM free Bup in U118MG (**Fig. 1B**), while in T98G, already a 20 µM **P-SS-Bup** with 81% decrease in metabolic activity elicited a statistically similar response to that of effective 2 µM free Bup (**Fig. 1C**). At 200 µM **P-SS-Bup**, all cell lines exhibited levels of metabolic suppression comparable to that achieved by 5 µM free Bup (**Fig. 1**), identifying it as a broadly effective concentration. The initial screening of SS-based Bup derivatives in 2D cultures provided the rationale for selecting effective concentrations for subsequent analyses. Given the well-documented increased resistance of GBM cells in 3D culture systems, 50 µM and 200 µM were selected as promising concentrations for further evaluation.

### 3.3 Modelling the release of Bup from polymer conjugate in the presence of GSH

Considering that the polymeric conjugate is substantially larger than either free Bup or the **SS-Bup** pro-drug, its intracellular efficacy is likely dependent on endocytosis-mediated uptake and subsequent intracellular drug release via GSH-triggered disulphide cleavage. To assess the model release kinetics of Bup from the polymer conjugates, additional experiments were conducted using a reductive environment containing 1 mM GSH at 25 °C over the course of 48 hours. The reaction progress was monitored by HPLC, allowing for the detection of several transient intermediates that gradually converted into free Bup.

As shown in **Figure 2**, both the rate and extent of Bup release were dependent on the initial concentration of the conjugate. There was no difference after 24 and 48 hours, approximately 39% of Bup was released at the 50 µM conjugate concentration, whereas only around 27% was liberated at 100 µM concentration. This inverse concentration-dependent release suggests that higher conjugate concentrations may hinder the efficiency of disulphide cleavage, possibly due to intermolecular interactions leading to formation of poorly soluble intermediates or limited accessibility of reducing agents within denser polymer matrices.

**Figure 2.**
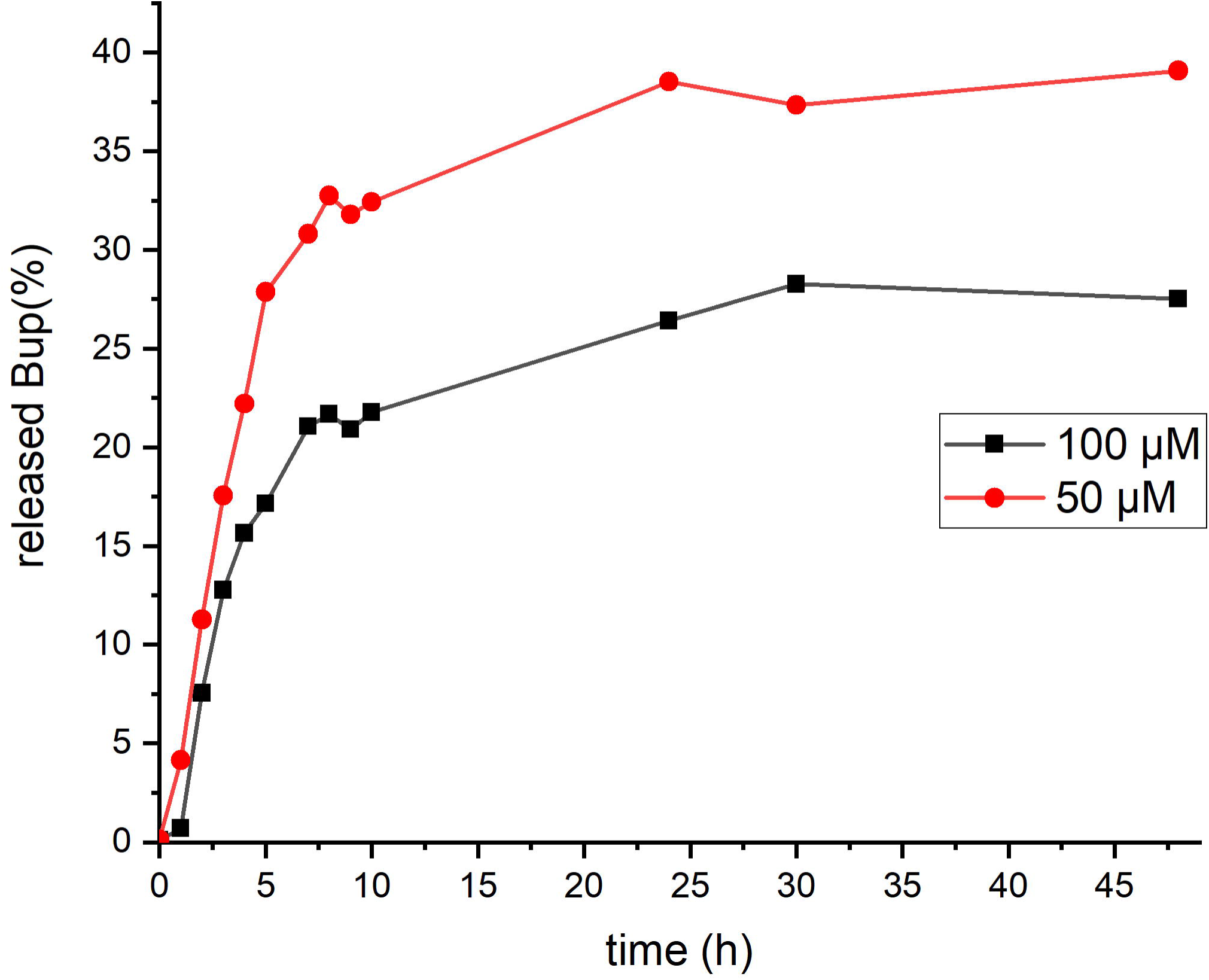
Release kinetics of Buparlisib from the polymer conjugate P-SS-Bup. Bup release from polymer conjugate **P-SS-Bup** in the presence of 1 mM GSH (25 °C, 75 mM tris-HCl buffer, pH 7.4). The release is expressed as a percentage relative to the complete cleavage with TCEP (100% reference).

Nevertheless, based on the observed dynamics, the actual amount of released Bup at higher conjugate concentrations can be estimated to be equal to or even higher than at lower concentrations. For example, 39% release from a 50 µM conjugate corresponds to 19.5 µM free Bup, while 27% release from 100 µM corresponds to 27 µM. Extrapolating this trend, we may speculate that approximately 15% release from a 200 µM conjugate could result in a similar effect to that of 30 µM of free Bup after 48 hours of treatment.

### 3.4 The synthesis of polymer conjugate with AP-Bup and the effect of drug modification on the metabolic activity of GBM cell lines in 2D conditions

For the synthesis of the **AP-Bup**, the precursor (**Prec1**) was first modified to introduce DBCO functional group, thus generating the second precursor **Prec2** with 4 mol% of reactive DBCO groups, as confirmed by UV/VIS determination of DBCO groups. The modification led to a slight increase in molecular weight (from 32.200 to 35.100 g/mol) and dispersity (Đ = 1.12). Importantly, both polymers retained low dispersity, supporting the preservation of the polymer backbone integrity throughout the synthetic steps. The reactive DBCO functionality in **Prec2** enables strain-promoted azide-alkyne cycloaddition (SPAAC), a bioorthogonal reaction suitable for further conjugation with azide-bearing derivatives.

The following polymer–drug conjugate **P-AP-Bup** was prepared by bioorthogonal conjugation of the azide-functionalised pro-drug **AP-Bup** with the DBCO-containing polymer precursor **Prec2** via SPAAC. This copper-free “click” reaction proceeded rapidly and efficiently in DMA at room temperature, as confirmed by HPLC analysis showing complete consumption of free **AP-Bup** within 1.5 hours. The resulting conjugate was purified by precipitation and gel filtration on Sephadex G-25, yielding the product in high purity and excellent yield (89 %) as a white powder. The final conjugate contained 10 wt% of Bup, which corresponds to the initial feed ratio of the **AP-Bup** (13.1 wt%) pro-drug in the reaction mixture. The use of SPAAC chemistry ensures high specificity and biocompatibility of the conjugation process, avoiding the need for metal catalysts or severe conditions.

To evaluate the effect of **AP-Bup** and its polymeric conjugate (**P-AP-Bup**) on GBM cells under 2D culture conditions, a 72-hour treatment was conducted using the pre-selected concentrations of 50 µM and 200 µM, as previously applied in the SS-based formulations.

In response to the **AP-Bup** treatment, we observed cell line-specific differences (**Fig. 3**), similar to the patterns noted for **SS-Bup**, where U118MG and T98G cells were more sensitive than U87MG. However, the overall inhibitory effect of the **AP-Bup** formulation was markedly lower. Treatment with 50 µM **AP-Bup** resulted in only a around 20% reduction in metabolic activity of U118MG and T98G cells, while only ≈5% decrease was observed in U87MG cells. Increasing the concentration to 200 µM led to a moderate decrease in metabolic activity in U87MG cells and approximately 50% reduction in U118MG (borderline insignificant inhibition compared to control; p_adj=0.0509; **Supplementary Table 1b**) and T98G cells (borderline significant inhibition compared to control, p_adj=0.0477, depicted in **Fig. 3C** as hatched bar; **Supplementary Table 1b**). Notably, the same concentration of **SS-Bup** elicited a substantially stronger inhibition across all three cell lines (**Fig. 1**).

**Figure 3.**
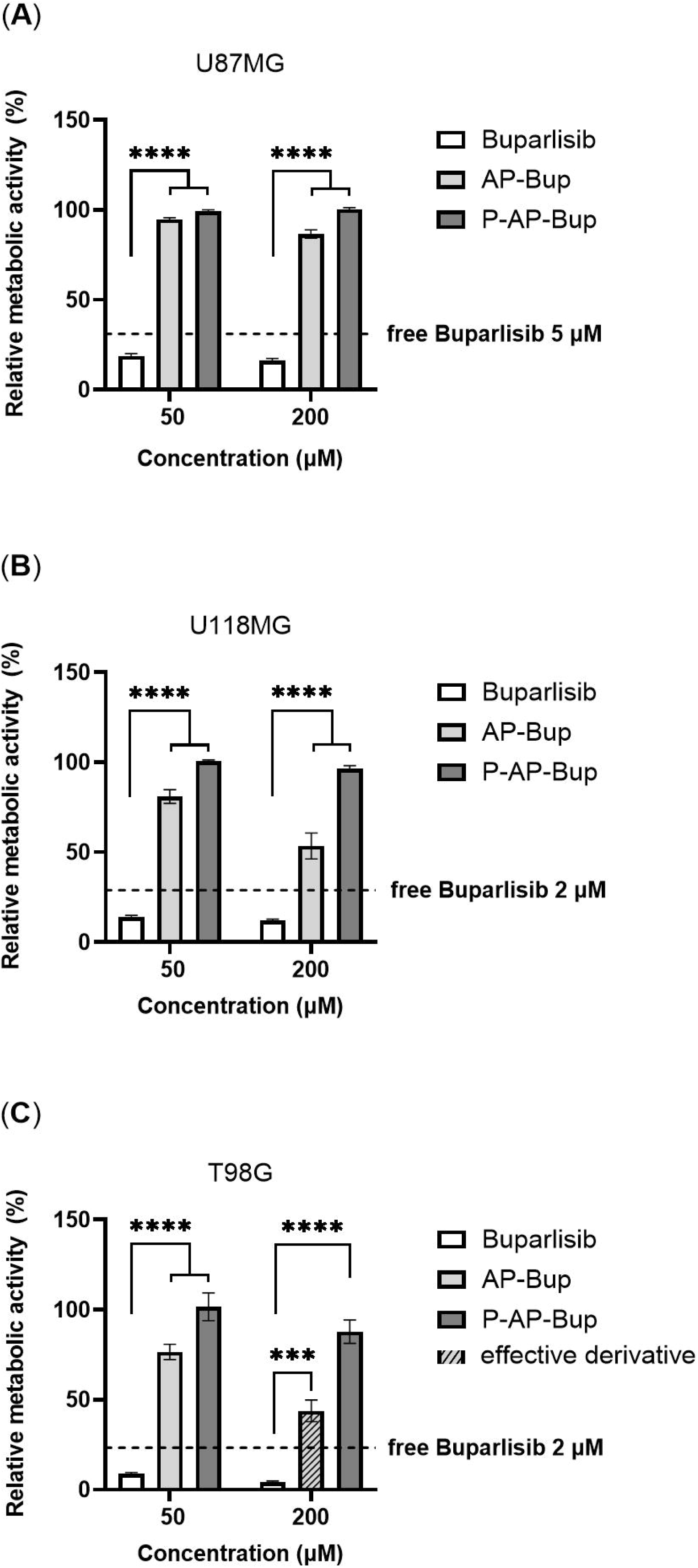
Effect of drug modification (AP) on the metabolic activity of GBM cell lines in 2D conditions. The effect of free Bup, Bup derivative containing an azide linker (**AP-Bup**), and its polymeric conjugated drug delivery construct (**P-AP-Bup**) on the metabolic activity of 2D-cultivated U87MG (**A**), U118MG (**B**), and T98G (**C**) cells. Values were normalised to the untreated control group (defined as 100 %). Data presented as mean ± SEM. Significance levels (adjusted p - values): [ns] (p_adj ≥ 0.05); [*] (p_adj < 0.05); [**] (p_adj < 0.01); [***] (p_adj < 0.001); [****] (p_adj < 0.0001). n=5-6

Treatment with the polymeric conjugate **P-AP-Bup** had a minimal impact on cell viability, with only a slight (around 10 %) reduction in metabolic activity observed in T98G cells (**Fig. 3C**). Collectively, these results indicate that the AP-based modification of Bup is substantially less effective than the SS-linked derivatives under the tested conditions. Since AP-based derivatives demonstrated minimal activity even at higher concentrations that were effective for SS-modified systems, these formulations were excluded from subsequent analysis.

### 3.5 Characterisation of the 3D spheroid models and the effect of drug modification on the GBM cell lines in 3D conditions

For further testing in 3D model of GBM, we generated spheroids from three GBM lines (U87MG, U118MG, and T98G). Monitoring of untreated spheroids revealed differences in spheroid growth dynamics, viability, and metabolic activity (**Fig. 4A, 4B, 4C**, respectively). The investigation of metabolic activity and cellular viability revealed a strong impairment in the case of the T98G cell line. Moreover, while the size of U87MG and U118MG spheroids increases during the cultivation, the T98G spheroids show a significant compaction. The pronounced loss of viability of T98G cells in 3D culture (**Fig. 4**), together with aberrant and reversed growth dynamics (**Fig. 4A**), renders the readouts unreliable and prevents meaningful analysis. Therefore, the T98G cell line was excluded from further analysis of treatment effects in 3D conditions.

**Figure 4.**
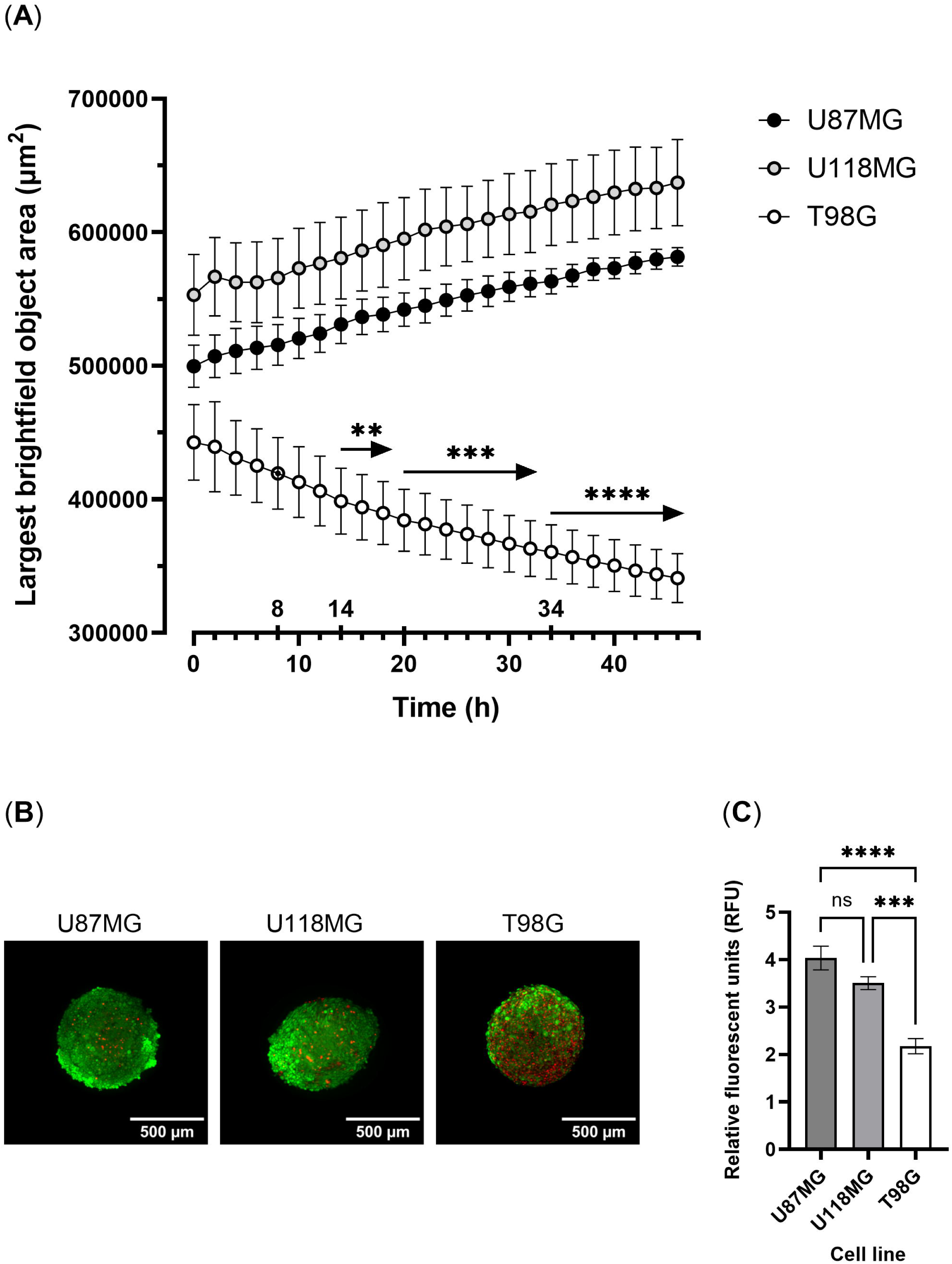
Growth dynamics, metabolic activity, and viability of 3D-cultured U87MG, U118MG, and T98G glioblastoma cell lines. (A) Spheroid size measured over 62 h using the Incucyte® Sx5 live-cell imaging system. Data presented as means ± SEM. Differences were analysed for each time point. First T98G timepoint value significantly different from both U87MG and U118MG values is depicted as a double circle. Significance levels (adjusted p - values): [**] (p_adj < 0.01); [***] (p_adj < 0.001); [****] (p_adj < 0.0001). n=6-8 (B) Live/dead staining of 3 days old U87MG, U118MG, and T98G spheroids using fluorescein diacetate (FDA) (live cells, green) and propidium iodide (PI) (dead cells, red); representative images from one biological replica. (C) Metabolic activity of spheroids measured after 3 days of culture. Data presented as mean ± SEM. Significance levels (adjusted p - values): [ns] (p_adj ≥ 0.05); [***] (p_adj < 0.001); [****] (p_adj < 0.0001). n=7

To evaluate the impact of Bup modification through pro-drug synthesis (**SS-Bup**) and subsequent conjugation to the HPMA polymer (**P-SS-Bup**), we performed a comparative analysis of their effects on 3D cultures of U87MG and U118MG cell lines. To further rationalise the selection of drug concentrations used in the subsequent 3D experiments, we additionally determined the IC50 values of free Bup, the **SS-Bup** derivative, and the polymeric conjugate **P-SS-Bup** in 2D cultures of U87MG and U118MG cells using a resazurin-based assay shown previously. In U87MG cells, the IC50 values were 2.482 µM for free Bup, 16.47 µM for **SS-Bup**, and 78.17 µM for **P-SS-Bup**, while in U118MG cells, the corresponding IC50 values were 0.6444 µM, 6.180 µM, and 32.77 µM, respectively (Supplementary Fig. 11). These results demonstrate a pronounced reduction in the apparent drug efficacy upon modification, supporting the use of higher concentrations in 3D culture systems. Based on these results, concentrations of 50 µM and 200 µM were selected for subsequent 3D experiments to ensure measurable biological effects across both cell lines.

Based on the observations from 2D experiments, where the concentrations of **P-SS-Bup** necessary to elicit an effect on metabolic activity comparable to that of lowest effective Bup were high (i.e. 200 µM in U87MG cells, and 50 µM in U118MG cells), and the IC50 calculation, such concentrations were selected for spheroid treatment. Spheroids were treated with the selected concentrations of substances, and subsequently monitored for the course of 62 hours (treatment period). Metabolic activity was assessed after the treatment period (**Fig. 5**).

**Figure 5.**
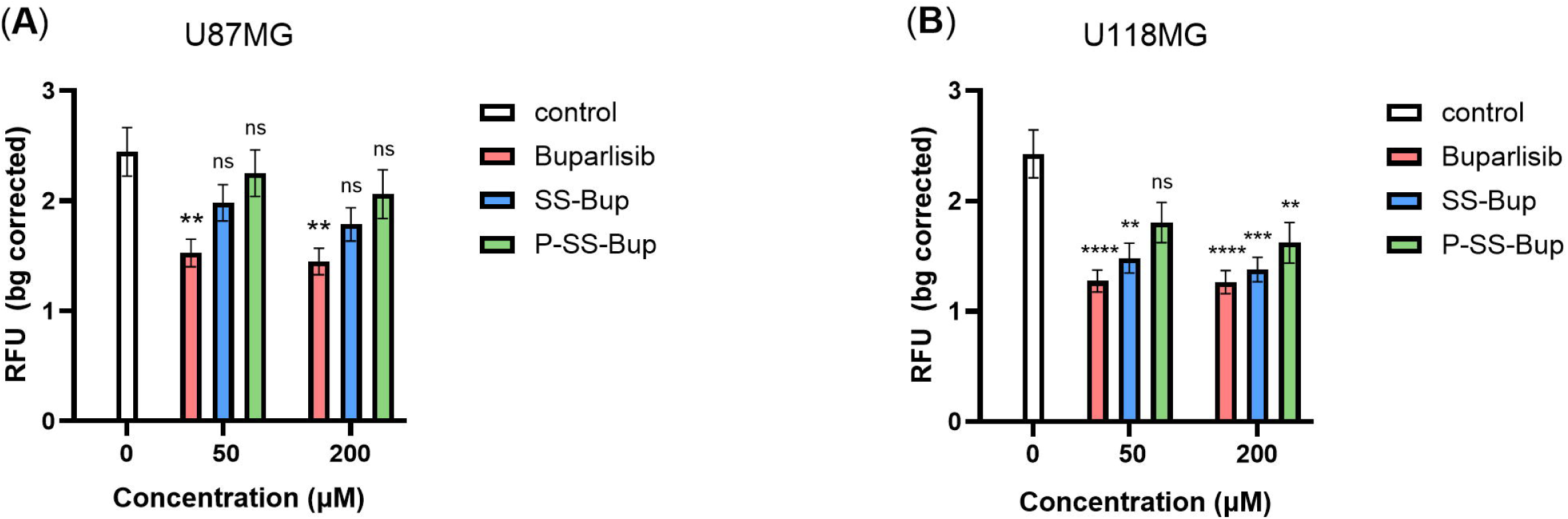
Effect of drug modification (SS) on the metabolic activity of GBM cell lines in 3D conditions. The effect of 72-hour treatment with free Bup red), pro-drug containing a disulphide bridge (**SS-Bup**; blue), and polymer-conjugated drug delivery construct (**P-SS-Bup**; green) in 50 µM and 200 µM concentrations on metabolic activity of 3D-cultured U87MG (**A**) and U118MG (**B**) glioblastoma cell lines. Data presented as mean relative fluorescence units (RFU) per group ± SEM. Significance levels (adjusted p - values): [ns] (p_adj ≥ 0.05); [**] (p_adj < 0.01); [***] (p_adj < 0.001); [****] (p_adj < 0.0001). n=9

In accordance with the results from 2D experiments (**Fig. 1A, 1B**), the modification of free Bup by introducing a disulphide (SS) linker/and polymeric backbone led to a reduction in its ability to inhibit the metabolic activity of GBM spheroids. In response to free Bup treatment, we observed a significant reduction in metabolic activity in both cell lines (**Fig. 5A, 5B**), while the treatment with derivatives was sufficient to cause a significant decrease in metabolic activity only in U118MG spheroids (**Fig. 5B**). Notably, in this cell line, the polymeric conjugate **P-SS-Bup** had a significant effect only at the highest tested concentration (200 µM), statistically indifferent from the other effective substances. The higher sensitivity of the U118MG cell line recapitulates the observations from 2D experiments.

Despite the seemingly insufficient inhibition of metabolic activity, especially in U87MG, both cell lines’ growth dynamics were significantly impaired in response to 62-hour treatment with both free Bup and its **SS-Bup** and **P-SS-Bup** derivatives (**Fig. 6A, 6B**). In drug-free conditions, spheroids from both cell lines exhibited continuous growth throughout the monitoring period, reaching approximately 120 % of the starting size by the end of the treatment period. Free Bup treatment induced an early and sustained impairment of growth dynamics. At the lower concentration (50 µM), spheroids sizes significantly diverged from the control within the first 10-12 hours, followed by a progressive reduction, ending below the starting size (≈85 %) after 62 hours. At the higher concentration (200 µM), the final reduction was less pronounced (≈95 %) (**Fig. 6A, 6B**).

**Figure 6.**
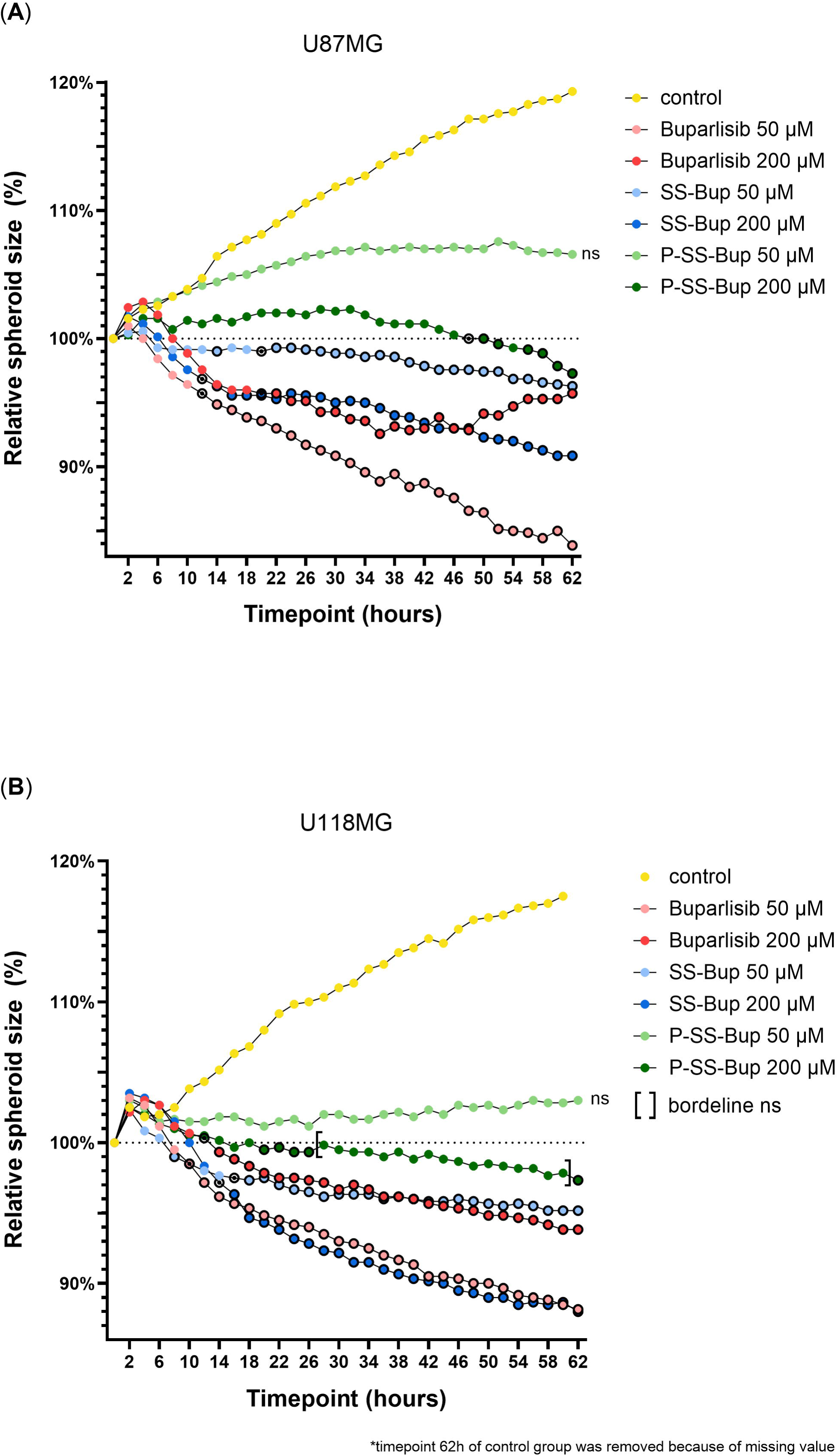
Effect of drug modification (SS) on the growth dynamics of GBM cell lines in 3D conditions. The effect of 72-hour treatment with free Bup (red), pro-drug containing a disulphide bridge (**SS-Bup**; blue), and polymer-conjugated drug delivery construct (**P-SS-Bup**; green) in 50 µM and 200 µM concentrations on growth dynamics of 3D-cultured U87MG (**A**) and U118MG (**B**) glioblastoma cell lines. Normalised data presented as means (SEM not shown for visual clarity). For each treated group, the first timepoint significantly different from control followed by continuously significant values is depicted as double circle. n=6-7

Both cell lines were also susceptible to growth impairment by the selected derivatives. In accordance with results from 2D resazurin assays (**Fig. 1A, 1B**), we observed a markedly delayed onset (12-20 hours for **SS-Bup**) of significant growth impairment (**Fig. 6A, 6B**). The treatment with **SS-Bup** resulted in a gradual decrease below the starting size, reaching around 95 % in response to the 50 µM dose. At 200 µM, both cell lines exhibited a more pronounced growth impairment, resulting in an endpoint size of 88-91 % of the starting size.

As expected, the polymeric conjugate **P-SS-Bup** had the least effect on spheroid size. Despite the statistically non-significant divergence from the untreated control at the lower concentration, we observed a clear stagnation of spheroid growth in both cell lines, with a delayed onset (30h timepoint) in the case of the U87MG spheroids. Moreover, the higher selected concentration resulted in a gradual decrease in the spheroid size. In response to 200 µM, we observed a growth stagnation and continuous significant divergence from the control, starting from the 48h timepoint in the case of the U87MG line, while the endpoint size (≈97 %) was comparable to that resulting from 50 µM **SS-Bup** and 200 µM free Bup treatment (≈96 %) (**Fig. 6A**). The impact of high concentration of **P**-**SS**-**Bup** on seemingly overall more sensitive U118MG spheroids was slightly above the statistically significant threshold between timepoints 28-60h (borderline ns, 0.0576 ≤ p_adj ≤ 0.0782), nevertheless, still resulted in a biologically meaningful gradual decrease below the starting size. Importantly, no significant effect of the empty, drug-free polymeric carrier on spheroid growth was observed at the same concentrations (Supplementary Fig. 12).

Overall, analysis of spheroid growth dynamics revealed that both GBM models were susceptible to treatment-induced growth impairment, even in cases where metabolic activity inhibition was limited or delayed. Statistically significant effects on growth dynamics were first observed in response to treatment with free Bup; with **SS-Bup**, they appeared with a delay, and the delay was greatest with the polymeric conjugate **P-SS-Bup**.

## 4 Discussion

This study establishes a versatile *in vitro* platform for a rational development and evaluation of polymer-based Bup delivery systems in glioblastoma, a malignancy with limited therapeutic options and high clinical urgency. Our approach integrated an advanced drug modification chemistry with both 2D and 3D *in vitro* models to identify reproducible and effective strategies that support the development of novel therapeutics and promote approaches aiming to reduce reliance on animal models.

We selected Bup, a pan-PI3K inhibitor, as a therapeutic agent and designed two strategies for its conjugation to pHPMA copolymers: one via strain-promoted azide–alkyne cycloaddition (SPAAC) introducing potentially enzymatically degradable carbamate linkage, and the other via a redox-sensitive disulphide linkage. At each stage of the drug modification process, we evaluated the cytotoxic efficacy of the derivatives and conjugates in three glioblastoma cell lines with distinct PTEN and p53 status, reflecting potential variability in treatment response (Lee et al., 2020, Schneiderman et al., 2015). Subsequently, the most promising derivatives and conjugates were tested in 3D spheroid models, which better mimic the *in vivo* tumour environment and are more physiologically relevant compared to 2D conditions. To our knowledge, this is the first report describing the selected Bup modifications and the first to evaluate their biological efficacy in 3D glioblastoma models.

Across all three GBM cell lines, free Bup exhibited potent suppressive effects on metabolic activity in a dose-dependent manner, achieving maximal inhibition at concentrations 2–5 µM. Comparable concentrations have been reported in other studies, showing low-µM activity, with variability across models (Chen et al., 2020, Netland et al., 2016, Bohnacker et al., 2017). These values reflect the variability in cellular sensitivity and emphasise the importance of selecting a representative cell line panel when conducting toxicological or pharmacological evaluations.

In agreement with previous reports, we observed cell line-specific differences in response to Bup. U118MG and T98G cells showed higher sensitivity to both free Bup and its derivatives compared to U87MG. Enhanced sensitivity of the T98G cells following Bup treatment has also been reported (Chen et al., 2020), however, in contrast to our findings, the authors observed similar levels of inhibition in U87MG cells. Another study reported higher Bup IC₅₀ for U87MG, compared to xenograft-derived GBM cells (Netland et al., 2016). In studies involving medulloblastoma cell lines, a similar pattern of cell line–dependent variability was observed, with Bup efficacy ranging across a broad spectrum depending on intrinsic features of the tested models (Zhao et al., 2017).

The differential response to Bup among GBM cell lines has frequently been attributed to variations in PTEN and p53 mutation status. While early studies proposed that low PTEN expression might enhance responsiveness to Bup due to amplified PI3K pathway activity, more recent evidence suggests a more complex relationship. In some cases, low PTEN levels have been associated with an increased resistance to Bup (Brandao et al., 2019, Cavazzoni et al., 2017). In opposite no consistent correlation between PTEN expression and Bup sensitivity was found across panels of GBM cell lines (Chen et al., 2020, Jane et al., 2014). Furthermore, both p53-dependent and p53-independent modes of Bup-induced cytotoxicity have been reported in other brain tumour models (Zhao et al., 2017), suggesting that the PI3K pathway may not be the sole determinant of therapeutic response.

In addition to PI3K inhibition, alternative mechanisms of Bup-induced cytotoxicity have been proposed. Bohnacker et al. demonstrated that at concentrations exceeding 1 µM, Bup induces cytotoxic effects that are independent of PI3K pathway inhibition. Specifically, they reported off-target inhibition of tubulin polymerisation, leading to microtubule destabilisation and cell cycle arrest. This PI3K-independent mechanism may contribute to the broad cytotoxicity of Bup at higher concentrations, even in cells with low PI3K signalling activity (Bohnacker et al., 2017).

Although our study was not aimed at elucidating the mechanisms underlying Bup cytotoxicity, the relatively high Bup concentrations used suggest that cytoskeletal destabilisation may have contributed significantly to the observed effects. Indeed, within the tested concentration range, we predominantly observed rapid cell detachment rather than gradual proliferation arrest, which is usually observed in response to PI3K inhibition, supporting the potential involvement of microtubule disruption.

Considering the modest clinical efficacy of Bup monotherapy at low concentrations (Wen et al., 2019) and its potential PI3K-independent cytotoxicity at higher doses (Bohnacker et al., 2017), optimising Bup-based therapy requires either combinatorial approaches with other therapeutics or the development of advanced drug delivery systems. Indeed, combining Bup with agents such as temozolomide, TH588, or AEW541 has been shown to improve treatment outcomes across various cancer models (Chen et al., 2020, Li et al., 2017, Anderson et al., 2015). In our study, we focused on the development of a drug delivery system designed to mitigate the off-target toxicity of Bup. Specifically, we engineered polymer–drug conjugates composed of an HPMA copolymer backbone, a cleavable linker, and the active Bup payload. Considering previous reports demonstrating that GSH levels are approximately fourfold higher in the tumour microenvironment compared to normal tissue (Xiong et al., 2021), the incorporation of a GSH-cleavable disulphide linker (SS-linker) into the drug delivery construct was considered a promising strategy. SS-based linkers have previously been employed in the development of polymeric conjugates with other chemotherapeutic agents, such as doxorubicin (Liang et al., 2018). To the best of our knowledge, however, no published studies have reported the application of this modification to Bup.

We found that the incorporation of a disulphide linker into the drug payload attenuated the suppressive effect of Bup across all tested GBM cell lines, while preserving the cell line-specific response profile observed with free Bup. The effective concentrations required to suppress metabolic activity were approximately 2.5- to 4-fold higher than those of free Bup. The modest increase in **SS-Bup** molecular weight is unlikely to impair passive diffusion. Instead, the lower efficacy of **SS-Bup** compared to its unmodified analogue most likely results from reduced intracellular drug availability, which depends on glutathione-mediated cleavage. Subsequent incorporation of the **SS-Bup** derivative into the larger **P**-**SS**-**Bup** polymeric construct resulted in a further shift of effective inhibitory concentrations toward higher doses. Despite this modulation, we observed a consistent, concentration-dependent reduction in metabolic activity across all tested cell lines, with maximal inhibition achieved within the 20–200 µM range, depending on the cell line. T98G cells remained the most sensitive to treatment, which could reflect differences in endocytic activity, intracellular trafficking, or glutathione availability that influence both the uptake and the release kinetics of the polymer-bound drug.

It is important to mention that this initial screening of effective concentrations was based on the resazurin assay, which reflects cumulative metabolic activity and is strongly influenced by cell number (Vieira-da-Silva and Castanho, 2023). Since the cells in control group continued to proliferate during the exposure period, the observed reduction in resazurin fluorescence indicated both the proliferation-inhibitory and the cytotoxic effects. Thus, the marked suppression of metabolic activity by 2 µM free Bup (and 5 µM in the case of the more resistant U87MG line) reflects effective inhibition of proliferation and onset of cell death. These values were therefore used as functional benchmarks for assessing the potency of Bup derivatives. Given that a) comparable inhibitory effects of derivatives (**SS-Bup** and **P**-**SS**-**Bup**) in 2D conditions were achieved only at substantially higher concentrations (being approximately 4–50 times higher, depending on the cell line), b) the IC50 calculation results confirmed this observation, and c) the fact that 3D cultured cells usually exhibit reduced drug sensitivity, concentrations of 50 µM and 200 µM were selected. These doses were the lowest at which SS-based formulations resulted in metabolic suppression equivalent to that of 2–5 µM free Bup, making them functionally comparable to the effective Bup concetrations in more treatment-resistant 3D models.

Replacing the disulphide linker with a SPAAC-based linker led to an apparent loss of activity of the polymeric conjugate (**P**-**AP**-**Bup**). However, this reduced effect is not due to the intrinsic inactivity of the polymeric system, but rather to the slow drug release. Even at the highest concentrations tested (50 µM and 200 µM), no significant inhibition was observed, and metabolic activity remained comparable to that of untreated controls. In contrast, the polymer-free derivative (**AP-Bup**) retained moderate activity at high concentrations, particularly in the U118MG and T98G cell lines. Following endocytic uptake, the linker is expected to be exposed to lysosomal esterases; nevertheless, the polymeric architecture of **P**-**AP**-**Bup** likely limits esterase accessibility and cleavage efficiency. Consequently, the active drug is released more slowly, resulting in reduced biological activity within the assay timeframe.

Although further investigation is necessary to elucidate the differences in cellular uptake and intracellular drug release between polymeric conjugates bearing distinct linker chemistries, we focused the subsequent experiments primarily on SS-based conjugates, excluding the AP-based formulations from further analysis.

The 3D *in vitro* behaviour of the three GBM cell lines differed markedly. While U87MG and U118MG cells formed spheroids that grew in size over time and maintained high viability and metabolic activity, T98G spheroids exhibited progressive shrinkage, accompanied by a decline in metabolic activity and a substantial loss of viability within three days of culture. Similar observations have been reported previously, where time-dependent decreases in T98G spheroid viability were noted in comparison to U87MG spheroids (Alves et al., 2023). However, in that study, the degree of shrinkage was less pronounced, potentially due to the presence of collagen in the 3D matrix (an element absent in our system). Given the compromised viability and metabolic activity of T98G spheroids prior to the drug treatment, this cell line was excluded from subsequent 3D drug testing experiments. Divergent behaviour of the T98G cell line in 3D conditions likely arises from its distinct genetic background (de Souza et al., 2022, Cesca et al., 2024) and high proliferation rate compared to the other cell lines used. We suggest that T98G may not be suitable for spheroid-based assays due to higher oxygen and nutrient demands stemming from these characteristics, which result in higher sensitivity to gradients in 3D conditions, ultimately reducing the cell viability in spheroids.

As expected, selected GBM cell lines cultured in 3D spheroids exhibited greater resistance to treatment compared to their 2D monolayer counterparts, as reported elsewhere (Musah-Eroje and Watson, 2019, Joseph et al., 2021, Rashidi et al., 2025, Lenin et al., 2021). This effect is generally not limited to a specific class of compounds, but rather reflects the structural and microenvironmental complexity of 3D systems. For example, U87MG spheroids were reported to be less sensitive to Erlotinib and Imatinib, requiring higher concentrations to achieve comparable responses (Rashidi et al., 2025). Similarly, screening of 65 compounds in patient-derived GBM models revealed substantial differences between 2D and 3D systems, as well as between individual samples, with only a limited number of compounds showing consistent efficacy across conditions (Lenin et al., 2021). Comparable results were also obtained in other models, where 3D-cultured cells showed increased resistance to agents such as Paclitaxel and Salinomycin analogues (Rafnsdottir et al., 2023), even under chemically defined, serum-free conditions. These observations support that 3D models represent a more stringent system for evaluating drug response.

Following 62 hours of treatment with free Bup, the reduction in metabolic activity in 3D was approximately twofold, whereas under 2D conditions, the same concentrations (50 and 200 µM) resulted in up to 90% inhibition. Notably, in U87MG spheroids, treatment with **SS-Bup** or **P**-**SS**-**Bup** had a minimal effect on metabolic activity. In contrast, U118MG spheroids remained relatively more sensitive to both derivatives, mirroring the response observed in 2D culture. It is important to emphasise that the overall metabolic activity of 3D-cultured cells was substantially lower than that of the 2D monolayers, consistent with previously reported findings (Duval et al., 2017, Guerrero-Lopez et al., 2025). Despite extending the resazurin incubation time to 4 hours to improve detection, we recognise that metabolic assays in 3D models may still underestimate drug effects or yield false-negative interpretations due to nonuniform dye penetration and reduced cellular metabolism.

Analysis of GBM spheroid growth dynamics during treatment, assessed by Incucyte® Sx5 imaging, revealed distinct patterns. In control drug-free conditions, both U87MG and U118MG spheroids showed an increase in diameter over 62 hours, indicating continued proliferation and 3D expansion. In contrast, nearly all treatment conditions suppressed spheroid growth, with some causing marked reductions in spheroid size. This was particularly evident in the free Bup and **SS-Bup** treatment groups. Although detailed mechanistic studies were beyond the scope of this work, we speculate that, similar to our 2D findings, exposure to high concentrations of free Bup or **SS-Bup** may disrupt cytoskeletal integrity. Such disruption could induce a shift toward a rounded cellular phenotype, leading to reduced intercellular spacing, increased packing density, and apparent spheroid compaction. Interestingly, the response to polymeric drug conjugates (**P**-**SS**-**Bup**) differed. While **P**-**SS**-**Bup** also inhibited spheroid growth compared to controls, it did not induce visible spheroid shrinkage. Instead, spheroid diameters remained relatively stable over time, suggesting growth arrest rather than compaction or cytotoxic collapse. The growth-suppressive effect of **P-SS-Bup** was most pronounced at 200 µM. At the same time, we may assume that the effect of **P-SS-Bup** on spheroid growth dynamics stems from Bup release from the conjugate rather than from a non-specific effect (e.g. mechanical obstruction) of the high polymer load (see Supplementary Fig. 12).

We presume that a combination of penetration rate, slower endocytic uptake of the polymer conjugates in comparison with the diffusion of the free Bup, together with the need for disulphide degradation and Bup activation, results in a higher efficacy at the periphery of the GBM spheroids and a reduced effect in the inner regions. Under these conditions, differences in delivery kinetics between freely diffusing small Bup molecules and release-limited **P-SS-Bup** can lead to distinct macroscopic phenotypes such as spheroid shrinkage versus suppression of spheroid expansion, as observed in our study.

In addition, the sustained but controlled activity of the polymeric construct aligns with the slow-release profile of Bup under reductive conditions and may better reflect the exposure in the context of solid tumours. In a reductive *in vitro* model with 1 mM GSH at 25 °C, mimicking cellular reductive conditions, complete linker cleavage from the polymer occurs; however, only about 40% of free Bup of the total amount bound in the conjugate is detected, probably due to the presence of other low-molecular-weight Bup intermediates. This supports delayed intracellular drug availability and reduced short-term potency and align with delayed PI3K inhibition confirmed by WesternBlot and immunodetection of downstream targets of PI3K in 2D conditions (**Supplementary Fig. 13**). These observations underscore the importance of integrating drug release kinetics into the interpretation of functional readouts in 3D cultures.

The **P-SS-Bup** seems to be an interesting candidate for further biological evaluation. The conjugate’s stability emphasises its potential to reduce side effects resulting from the release and early activation of free drug in the bloodstream. The observed reduction in the *in vitro* potency of the conjugated Bup, compared to free drug, should be interpreted in the context of its delivery-oriented design and release-dependent mode of action, while reductive degradation in the cytosol of tumour cells supports its suitability for controlled intracellular delivery in glioblastoma treatment strategies.

Although the present study was not designed to directly compare the mechanisms of action of the unconjugated and polymer-conjugated Bup, but rather to provide a rationale for the development of controlled drug-delivery systems, this represents a limitation of the current work. However, we may presume that the growth inhibition observed in 3D spheroids is attributed to, alas not exclusively, to the PI3K pathway blockade (as confirmed for U87MG cells in 2D conditions in **Supplementary Fig. 13**), and may also involve PI3K-independent off-target activity of Bup, such as cytoskeletal effects that become more pronounced depending on the effective exposure level.

Further mechanistic studies are required for the direct assessment of PI3K pathway engagement in spheroids. Measurement of downstream signalling markers such as phosphorylated AKT, S6, or 4EBP1 under matched exposure conditions would provide evidence of PI3K pathway inhibition in the 3D setting. In parallel, analysis of mitotic markers (e.g., phospho-histone H3) or microtubule organization could help to determine whether microtubule-related mechanisms contribute to the observed growth inhibition and clarify the relative contribution of different mechanisms in the activity of polymer-based Bup delivery systems.

At the same time, the *in vitro* activity of Bup derivatives was evaluated using Bup-equivalent concentrations to enable direct comparison with the free drug. However, this approach does not fully reflect the *in vivo* behaviour of polymer–drug conjugates, including their biodistribution and release kinetics. In this context, the effective dosing will ultimately depend on systemic exposure, tumour accumulation, and controlled drug release. In future *in vivo* studies, integrated pharmacokinetic and pharmacodynamic analyses will be required to define the relationship between administered dose, tumour exposure, and pathway inhibition. Such studies will be essential to establish dosing strategies for this and related polymer-based drug delivery systems, particularly when additional modifications (e.g. targeting moieties) are introduced that may further alter drug availability and efficacy.

## 5 Conclusion

By systematically evaluating Bup and its polymeric derivatives across different cell lines, linker chemistries, concentrations, and culture conditions, our study provides a comparative framework for rational drug delivery design in glioblastoma. Although mechanistic insights into cellular uptake and intracellular release remain to be fully elucidated, our findings offer critical guidance for selecting optimal linker strategies, effective concentration ranges, and responsive 3D cellular models, thereby establishing a reproducible *in vitro* platform that supports the continuous development of polymer–drug conjugates.

Our findings additionally underscore that conventional 2D models are insufficient for a comprehensive evaluation of therapeutic efficacy in solid tumours such as GBM. 3D *in vitro* models represent a key class of New Approach Methodologies (NAMs), as they provide a higher level of biological relevance. Importantly, when appropriately optimised and combined with complementary readouts, 3D models have the potential to enhance the predictive value of preclinical drug screening. An enhanced translational relevance, together with the development and use of new xeno-free cultivation strategies (Rafnsdottir et al., 2023) may in turn reduce the reliance on animal models and animal-derived products by enabling more informed compound prioritisation and early elimination of ineffective candidates in xeno-free settings, thereby contributing to the implementation of the 3Rs principles in preclinical testing.

Overall, this work goes beyond classical drug testing to establish a rational workflow for preclinical evaluation of polymeric nanomedicines, providing critical insights into the relationships between chemical design, biological context, and model relevance. The results lay a foundation for further refinement of Bup-based delivery systems and support the broader transition toward mechanism-informed, animal-free approaches in treatment development.

## 7 Author Contributions

JH, YP, TE, RP and PJ – concept and design of the study, data analysis and interpretation; JH, YP, RP –manuscript writing; JH,YP, AS, KP, DM, MP, MS, RP - experiments execution, collection and assembly of data; JH, YP, KP, DM – *in vitro* modelling and cell-based assays; AS, MP, MS, RP – synthesis and characterisation of polymeric conjugates; TE, RP, PJ - critical review of the manuscript, administrative and financial support; RP and PJ - supervision of the study. All authors read and approved the final manuscript.

## 9 Acknowledgments

The authors acknowledge Imaging Methods Core Facility at BIOCEV, institution supported by the MEYS CR (LM2023050 Czech-BioImaging) for their support & assistance in this work. Specifically, RNDr. Zuzana Čočková, Ph.D. for assistance with data analysis, and Ing. Dalibor Pánek, Ph.D. for support with confocal microscope image acquisition.

## 11 Supplementary Material

- -**Supplementary Material 1**

- **Supplementary Table 1.** Statistical analysis of effectiveness of Bup and its derivatives in comparison to control.
- -**Supplementary Material 2**

- **Supplementary Figure 1**. Synthesis of **AP-Bup** derivative scheme.
- **Supplementary Figure 2**. Mass spectrum of **AP-Bup.**
- **Supplementary Figure 3**. Synthesis of **SS-Bup** derivative.scheme.
- **Supplementary Figure 4**. NMR spectra of Ma-b-Ala-OH, Ma-b-Ala-TT and polymer precursor **Prec1**.
- **Supplementary Figure 5.** SEC chromatograms of **Prec1** (**A**) and **P-SS-Bup** (**B**).
- **Supplementary Figure 6**. Synthesis of polymer conjugate **P-SS-Bup** scheme (**A**) and HPLC chromatograms of the conjugation reaction (**B**).
- **Supplementary Figure 7**. Synthesis of polymer conjugate **P-AP-Bup** scheme.
- **Supplementary Figure 8**. Determination of 100% Buparlisib release using tris(2-carboxyethyl)phosphine (TCEP, 1 mM) under identical conditions (75 mM Tris-HCl, pH 7.4, 25 °C).
- **Supplementary Figure 9**. Timeline of the treatment regimes for 2D (**A**) and 3D (**B**) experiments. Created in BioRender. Havelková, J. (2026) https://BioRender.com/ee01iwb(licensed under CC BY 4.0.).
- **Supplementary Figure 10.** Effect of solvent DMSO on metabolic activity of cells in 2D conditions (**A**) and spheroid growth dynamics in the 3D conditions (**B, C**).
- **Supplementary Figure 11.** IC50 charts and calculation for Bup, **SS-Bup** and **P-SS-Bup** in U87MG (**A**) and U118MG (**B**) cell lines.
- **Supplementary Figure 12.** Effect of conjugate precursor **Prec1** on U87MG (**A**) and U118MG (**B**) spheroids growth dynamics.
- **Supplementary Figure 13.** Effect of Bup and **P-SS-Bup** on the level of phosphorylated AKT protein (p-AKT) in U87MG cells under 2D conditions.
- -**Supplementary Material 3**. SEM values omitted in growth dynamics charts (**Fig. 6A, 6B**).

## 12 Data Availability Statement

The datasets for this study can be found in the Zenodo at [https://doi.org/10.5281/zenodo.19261011] The repository includes all figures and raw and processed data used in this research.

