## Supplementary material for "Modular *in vitro* evaluation of Buparlisib-polymeric nanomedicines in 2D and 3D models of glioblastoma": Supplementary Material 1.docx

Jarmila Havelková^1,2,3*^, Yuriy Petrenko^1,2^, Anna Stehlíková^4^, Dana Mareková^1^, Kateřina Pešková^1^, Michal Pechar^4^, Martin Studenovský^4^, Tomáš Etrych^4^, Robert Pola^4*^, Pavla Jendelová^1^

^1^Department of Neuroregeneration, Institute of Experimental Medicine, Czech Academy of Sciences, Videňská 1083, 14200, Prague, Czech Republic

^2^Laboratory of Biomaterials and Tissue Engineering, Institute of Physiology, Czech Academy of Sciences, Videňská 1083, 14200, Prague, Czech Republic

^3^Faculty of Science, Charles University, Albertov 6, 12800, Prague, Czech Republic

^4^Department of Biomedical Polymers, Institute of Macromolecular Chemistry, Czech Academy of Sciences, Heyrovského nám. 2, 16200, Prague, Czech Republic

*** Correspondence:**
M.Sc. Jarmila Havelková, Ing. Robert Pola, Ph.D.


### 1 Supplementary Material 1

**Supplementary Table 1**. Statistical analysis of the effectiveness of Bup and its derivatives in comparison to control.

a)

|  | **Cell line** | | | | | |
| --- | --- | --- | --- | --- | --- | --- |
|  | **U87MG** | | **U118MG** | | **T98G** | |
| **Control vs:** | Summary | p_adj | Summary | p_adj | Summary | p_adj |
| **Bup 2µM** | ns | 0.3147 | *** | 0.0002 | *** | 0.0003 |
| **Bup 5µM** | *** | 0.0002 | **** | <0.0001 | **** | <0.0001 |
| **SS‑Bup 2µM** | ns | >0.9999 | ns | 0.9861 | ns | >0.9999 |
| **SS‑Bup 5µM** | ns | >0.9999 | ns | 0.0849 | *** | 0.0006 |
| **SS‑Bup 20µM** | ** | 0.0071 | *** | 0.0002 | **** | <0.0001 |
| **SS‑Bup 50µM** | *** | 0.0001 | **** | <0.0001 | **** | <0.0001 |

*Welsch’s ANOVA and Dunnett's T3 multiple comparisons post-hoc test

b)

|  | **Cell line** | | | | | |
| --- | --- | --- | --- | --- | --- | --- |
|  | **U87MG** | | **U118MG** | | **T98G** | |
| **Control vs:** | Summary | p_adj | Summary | p_adj | Summary | p_adj |
| **Bup 2µM** | ns | 0.9645 | * | 0.0118 | *** | 0.0003 |
| **Bup 5µM** | * | 0.0119 | ** | 0.0023 | *** | 0.0003 |
| **Bup 50µM** | ** | 0.0046 | ** | 0.0029 | *** | 0.0002 |
| **Bup 200µM** | ** | 0.0038 | ** | 0.0026 | *** | 0.0002 |
| **AP‑Bup 50µM** | ns | >0.9999 | ns | 0.7478 | ns | 0.6434 |
| **AP‑Bup 200µM** | ns | 0.9396 | ns | 0.0509 | * | 0.0477 |
| **P‑AP‑Bup 50µM** | ns | >0.9999 | ns | >0.9999 | ns | >0.9999 |
| **P‑AP‑Bup 200µM** | ns | >0.9999 | ns | >0.9999 | ns | 0.9714 |

*Welsch’s ANOVA and Dunnett's T3 multiple comparisons post-hoc test
