## Supplementary material for "Modular *in vitro* evaluation of Buparlisib-polymeric nanomedicines in 2D and 3D models of glioblastoma": Supplementary Material 2.docx

Jarmila Havelková^1,2,3*^, Yuriy Petrenko^1,2^, Anna Stehlíková^4^, Dana Mareková^1^, Kateřina Pešková^1^, Michal Pechar^4^, Martin Studenovský^4^, Tomáš Etrych^4^, Robert Pola^4*^, Pavla Jendelová^1^

^1^Department of Neuroregeneration, Institute of Experimental Medicine, Czech Academy of Sciences, Videňská 1083, 14200, Prague, Czech Republic

^2^Laboratory of Biomaterials and Tissue Engineering, Institute of Physiology, Czech Academy of Sciences, Videňská 1083, 14200, Prague, Czech Republic

^3^Faculty of Science, Charles University, Albertov 6, 12800, Prague, Czech Republic

^4^Department of Biomedical Polymers, Institute of Macromolecular Chemistry, Czech Academy of Sciences, Heyrovského nám. 2, 16200, Prague, Czech Republic

*** Correspondence:**
M.Sc. Jarmila Havelková, Ing. Robert Pola, Ph.D.


### 1 Supplementary Material 2

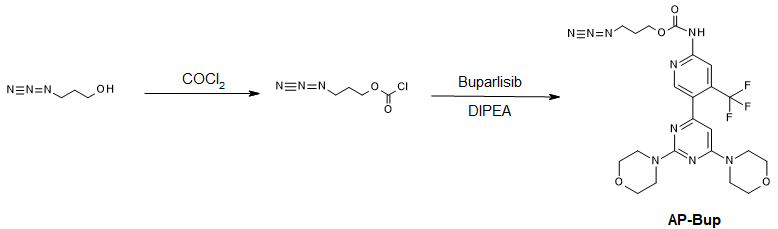

**Supplementary Figure 1.** Scheme of the synthesis of **AP-Bup** derivative.

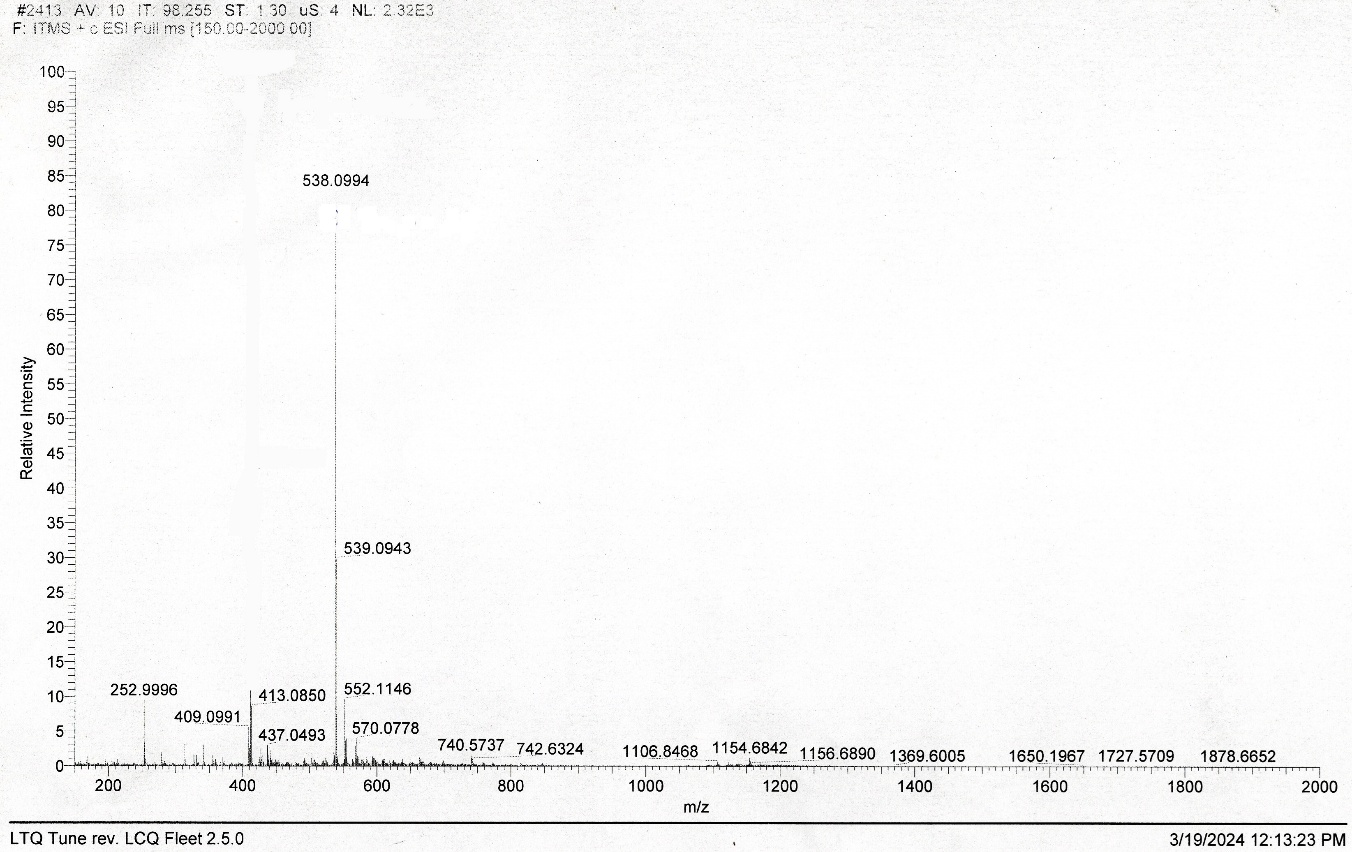

**Supplementary Figure 2.** Mass spectrum of **AP-Bup** ([M+H]⁺ = 538.0944)performed on an LCQ Fleet mass analyser with electrospray ionisation (ESI-MS).

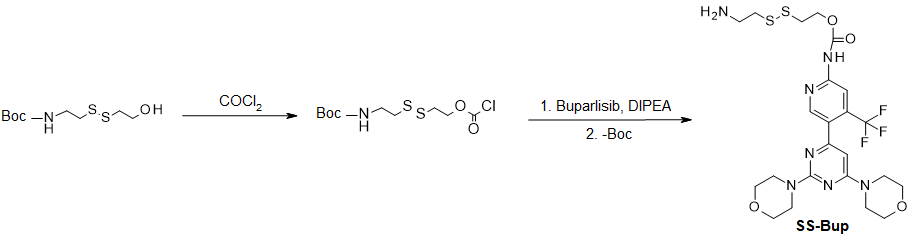

**Supplementary Figure 3.** Scheme of the synthesis of **SS‑Bup** derivative.

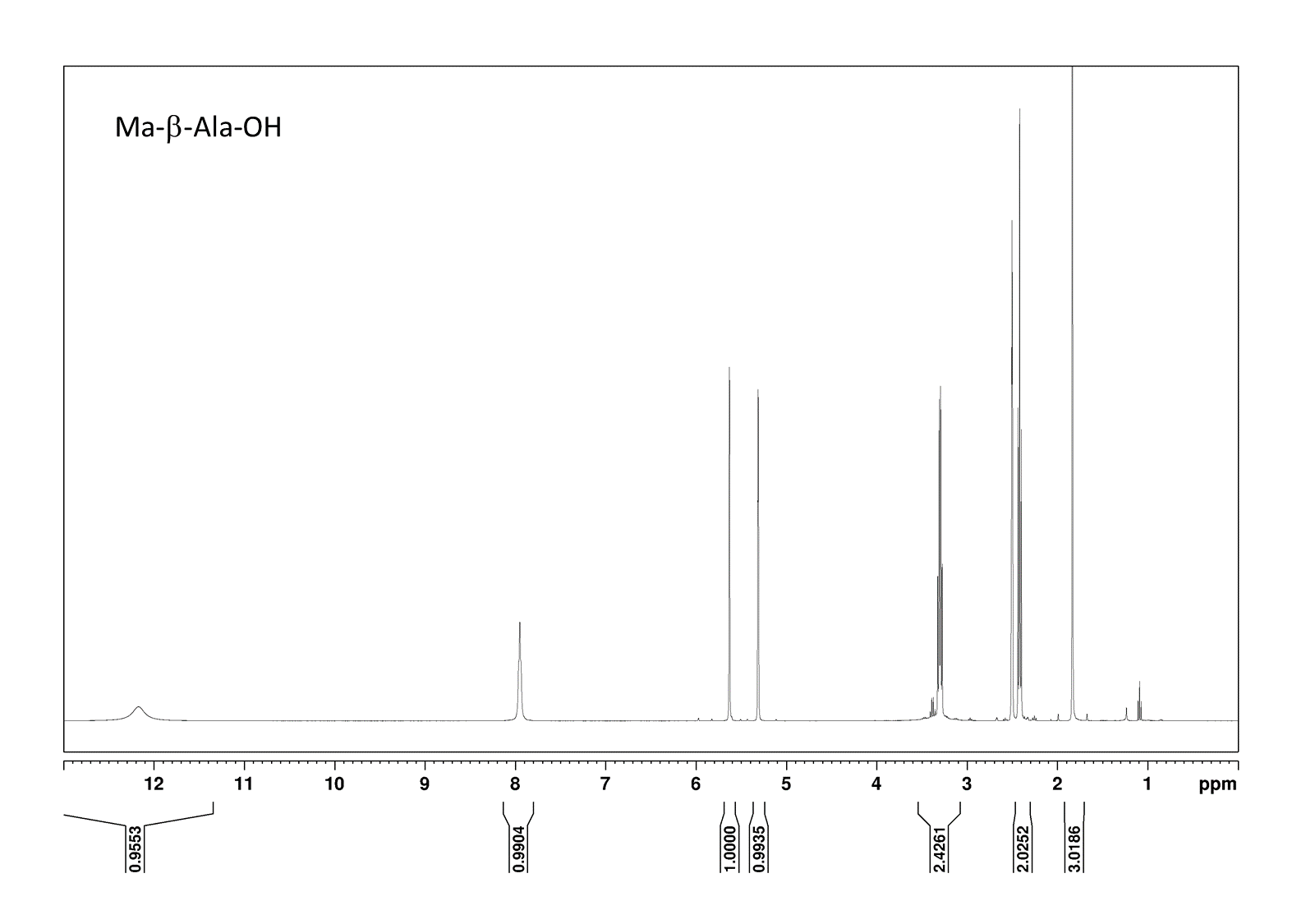

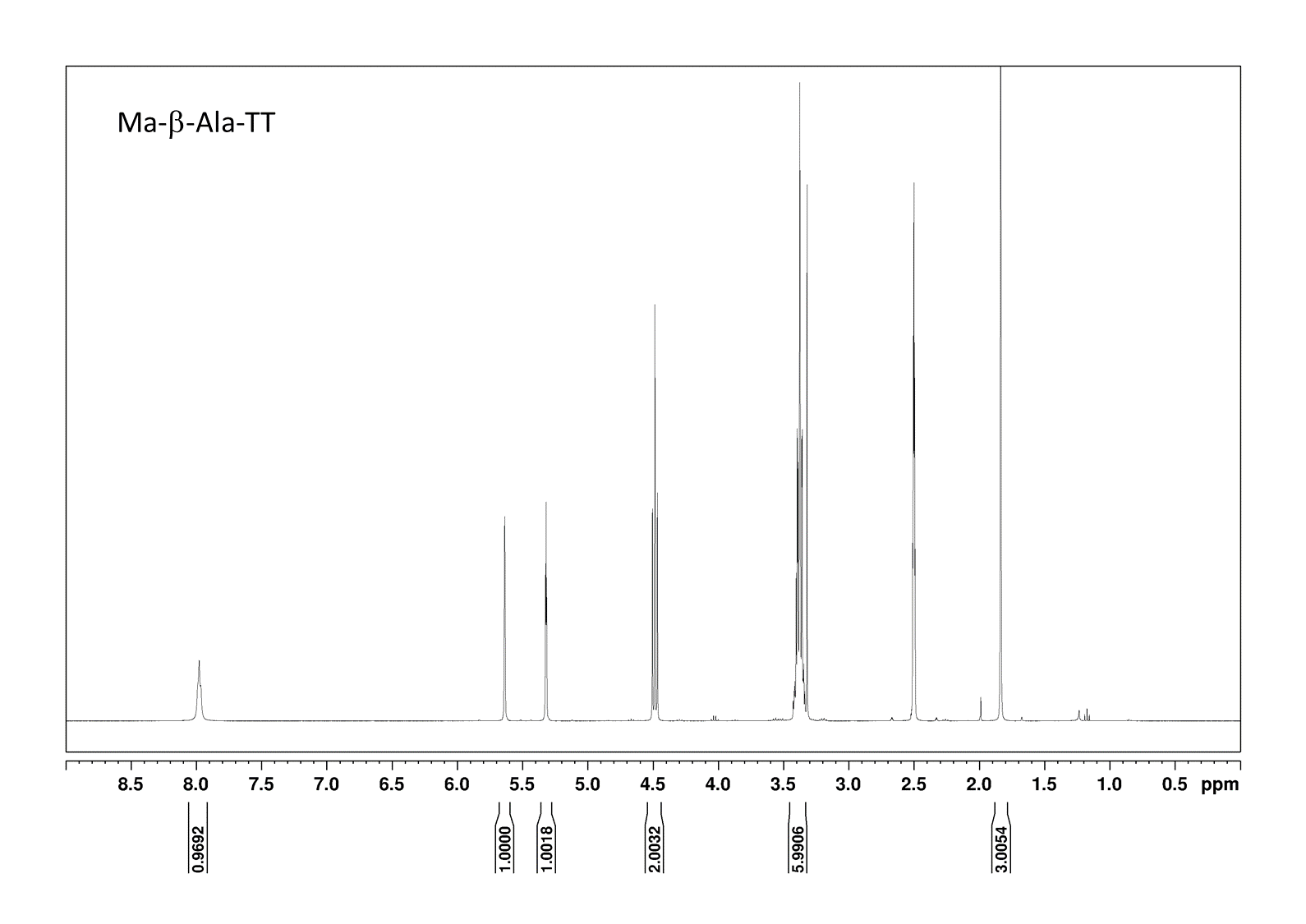

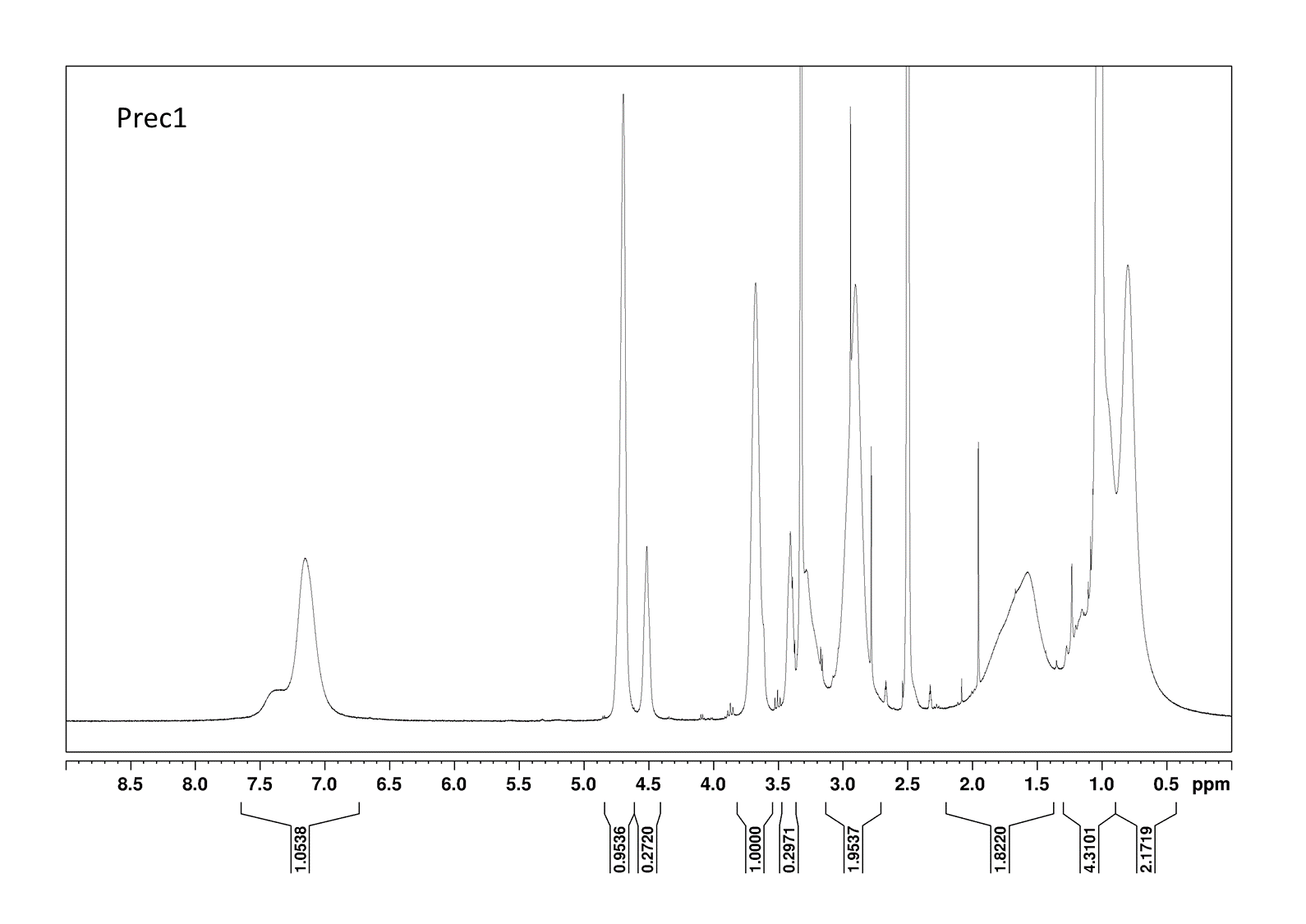

**Supplementary Figure 4.** NMR spectra of Ma-β-Ala-OH, Ma-β-Ala-TT and polymer precursor **Prec1**.

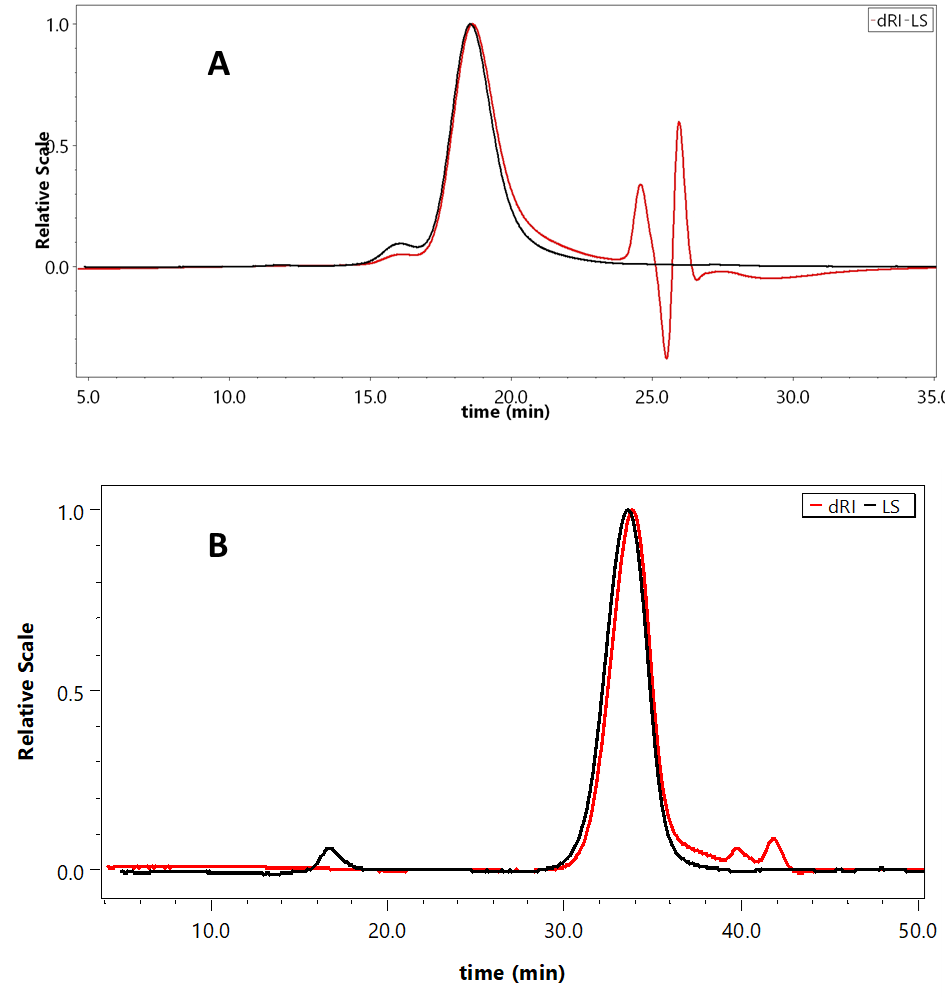

**Supplementary Figure 5.** SEC chromatograms of **Prec1** (MW= 32 200 g/mol) measured on a TSKgel G3000SWXL column (A) and **P-SS-Bup** (MW= 54 00 g/mol) measured on a Superose 6 Increase 10/300 GL column (B).

**A**

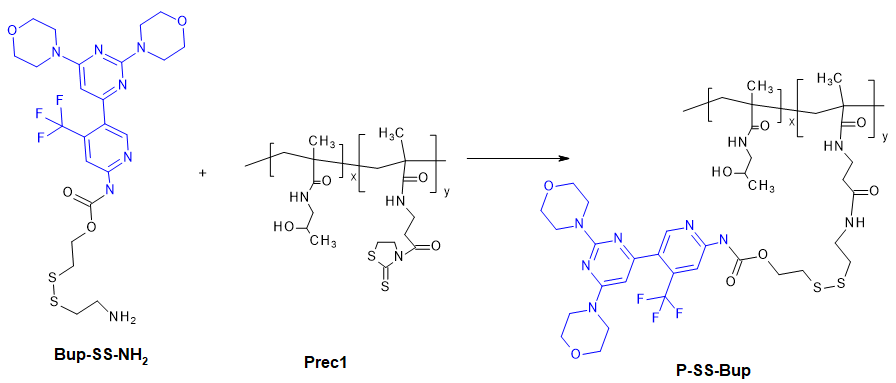

**B**

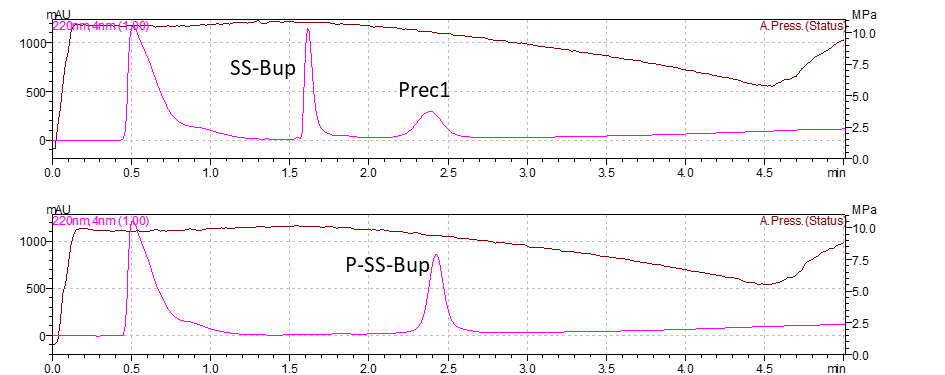

**Supplementary Figure 6.** Scheme of the synthesis of polymer conjugate **P‑SS‑Bup** (A) and HPLC chromatograms (220 nm) of the conjugation reaction (B).

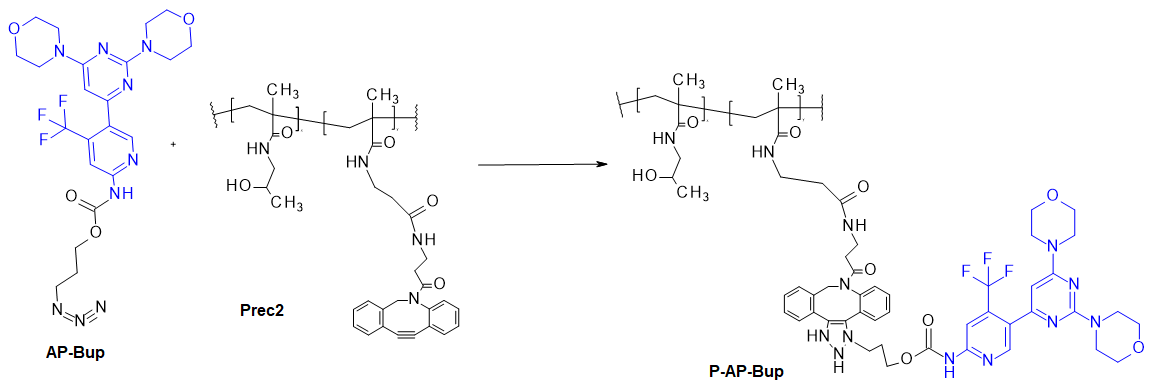

**Supplementary Figure 7.** Scheme of the synthesis of polymer conjugate **P‑AP‑Bup**.

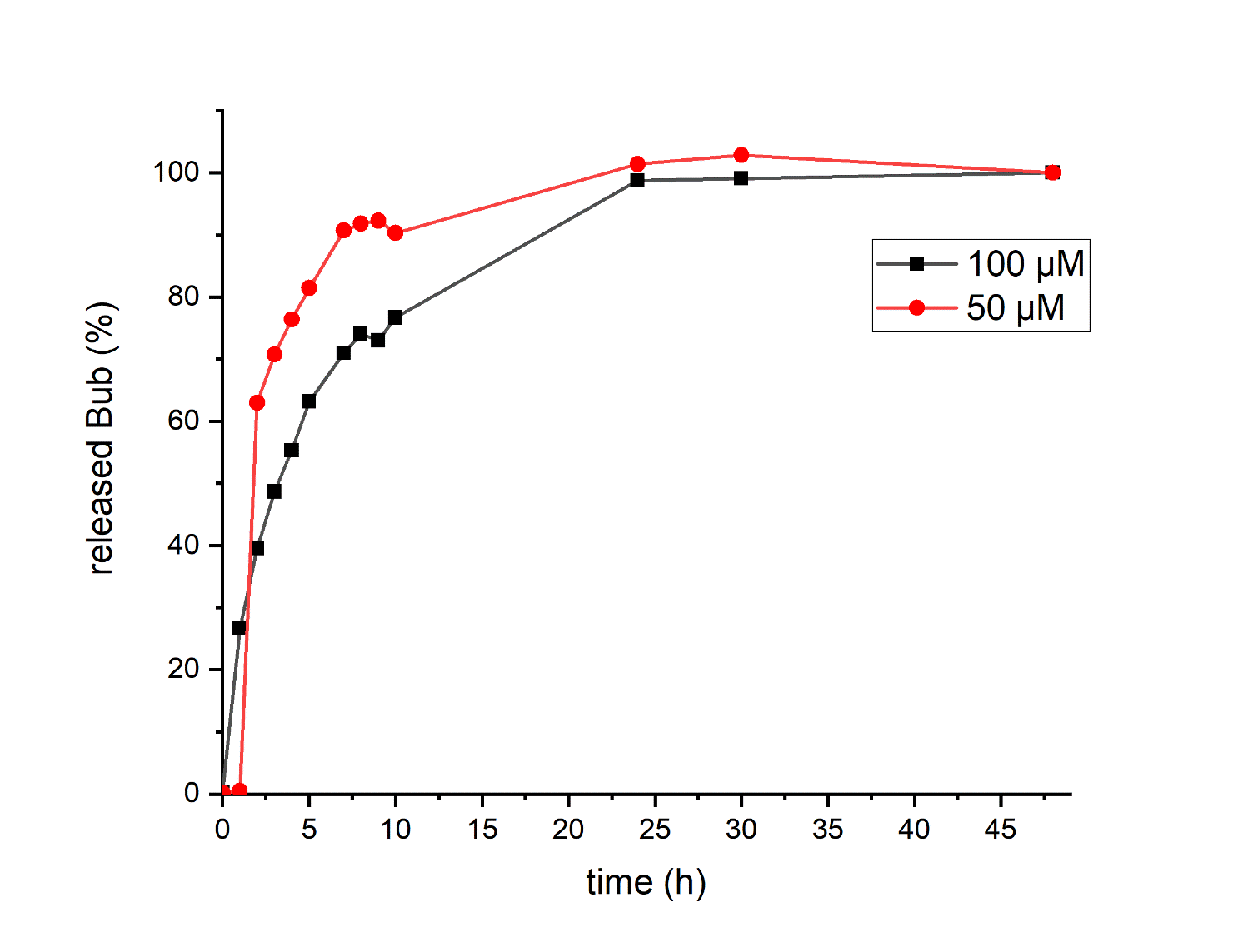

Supplementary Figure 8. Determination of 100% Buparlisib release using tris(2-carboxyethyl)phosphine (TCEP, 1 mM) under identical conditions (75 mM Tris‑HCl, pH 7.4, 25 °C). The experiment confirmed complete cleavage of the disulphide linkage and was used to establish the reference peak area for calculation of relative Buparlisib release in GSH‑mediated assays.

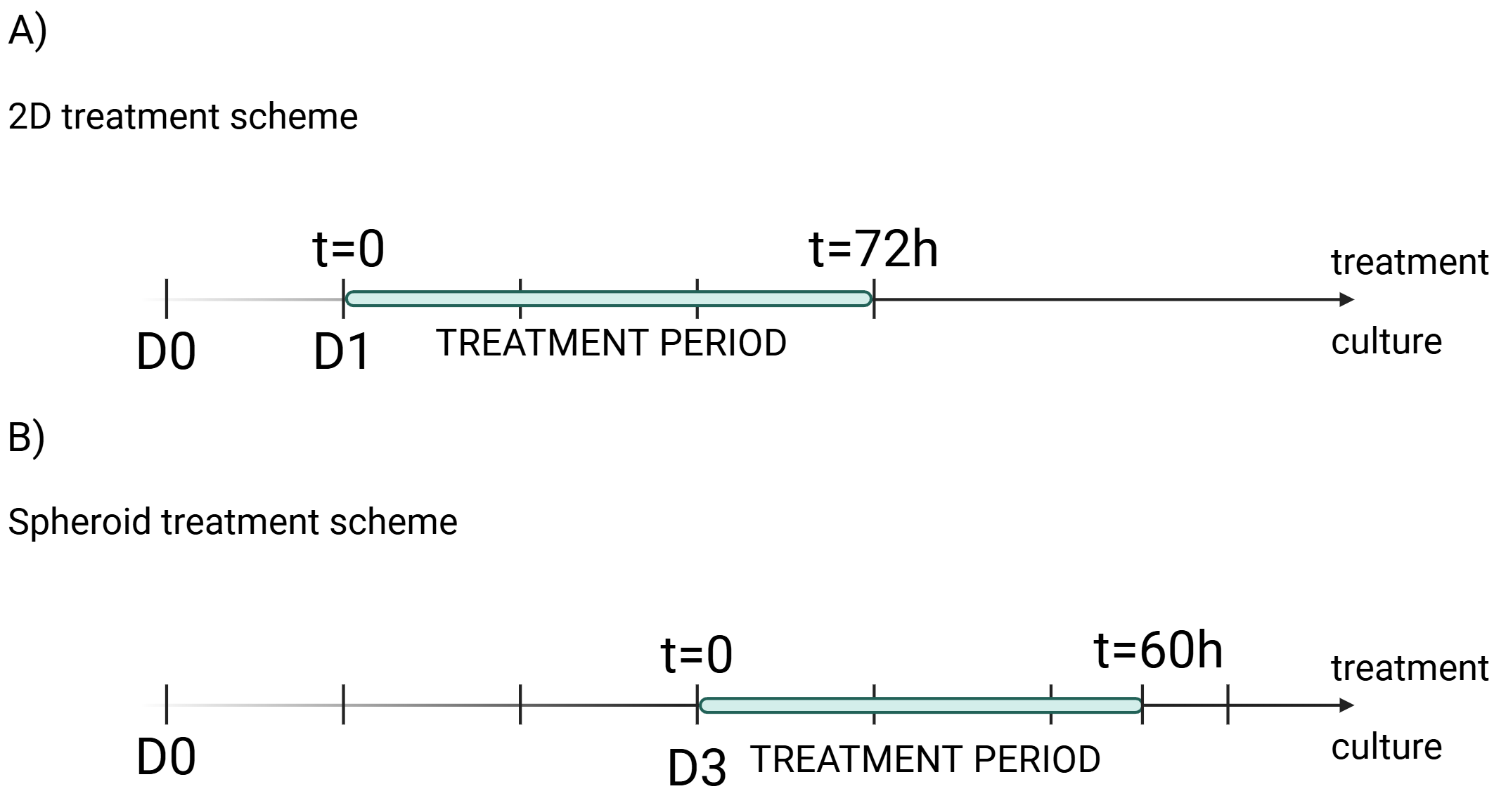

**Supplementary Figure 9.** Timeline of the treatment regimes for 2D (**A**) and 3D (**B**) experiments. Created in BioRender. Havelková, J. (2026) <https://BioRender.com/ee01iwb> (licensed under CC BY 4.0.).

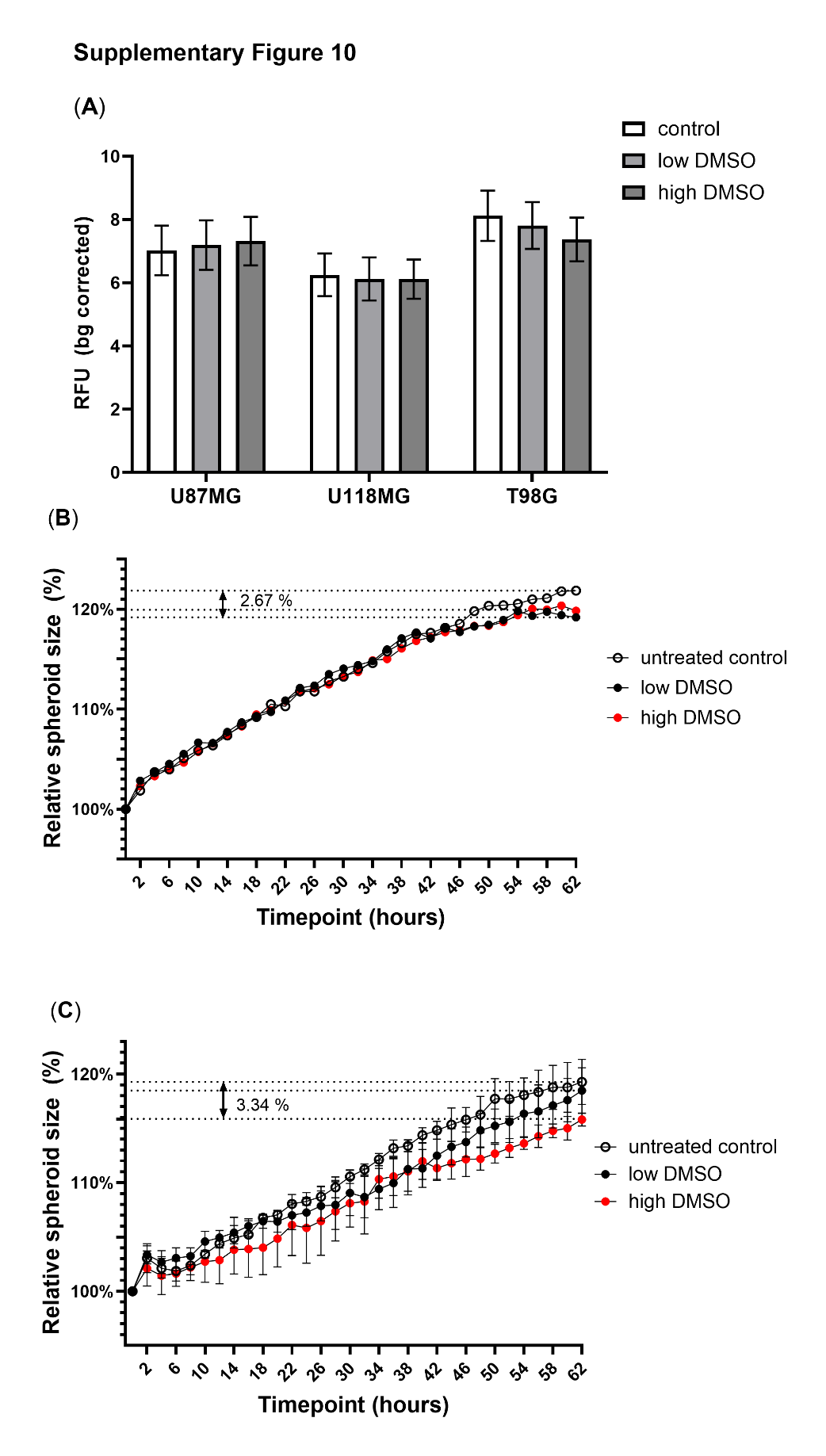

**Supplementary Figure 10.** Effect of solvent (DMSO) presence in cultivation media on metabolic activity of U87MG, U118MG, and T98G cells in 2D conditions (A; raw data plotted; n=6) and on U87MG and U118MG spheroid growth dynamics (B, C; data normalized to t=0 plotted; n=3).

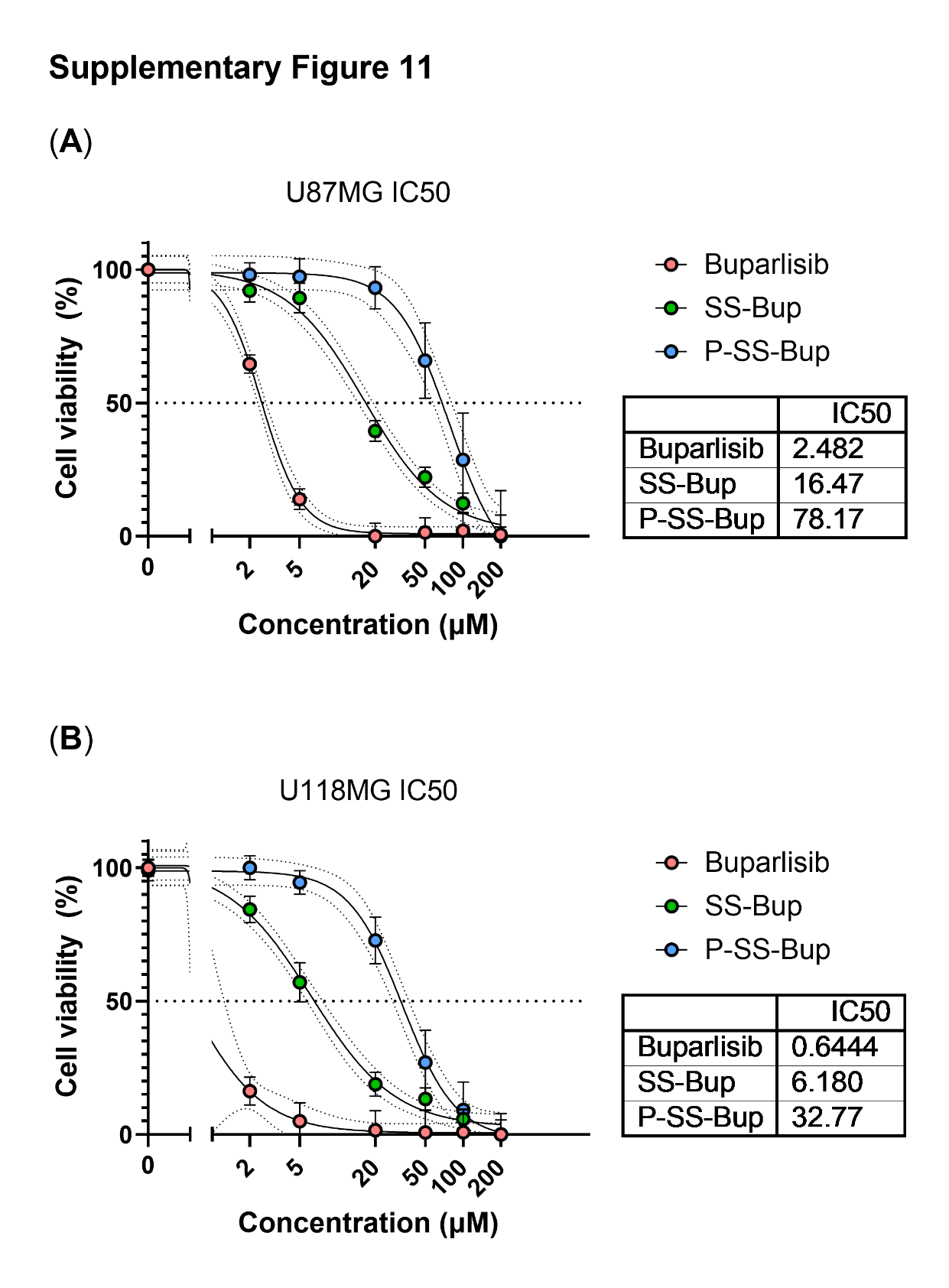

**Supplementary Figure 11.** IC50 charts and calculation for Bup, **SS-Bup**, and **P-SS-Bup** exposure for 72 hours in U87MG (A) and U118MG (B) cells in 2D conditions. n=4

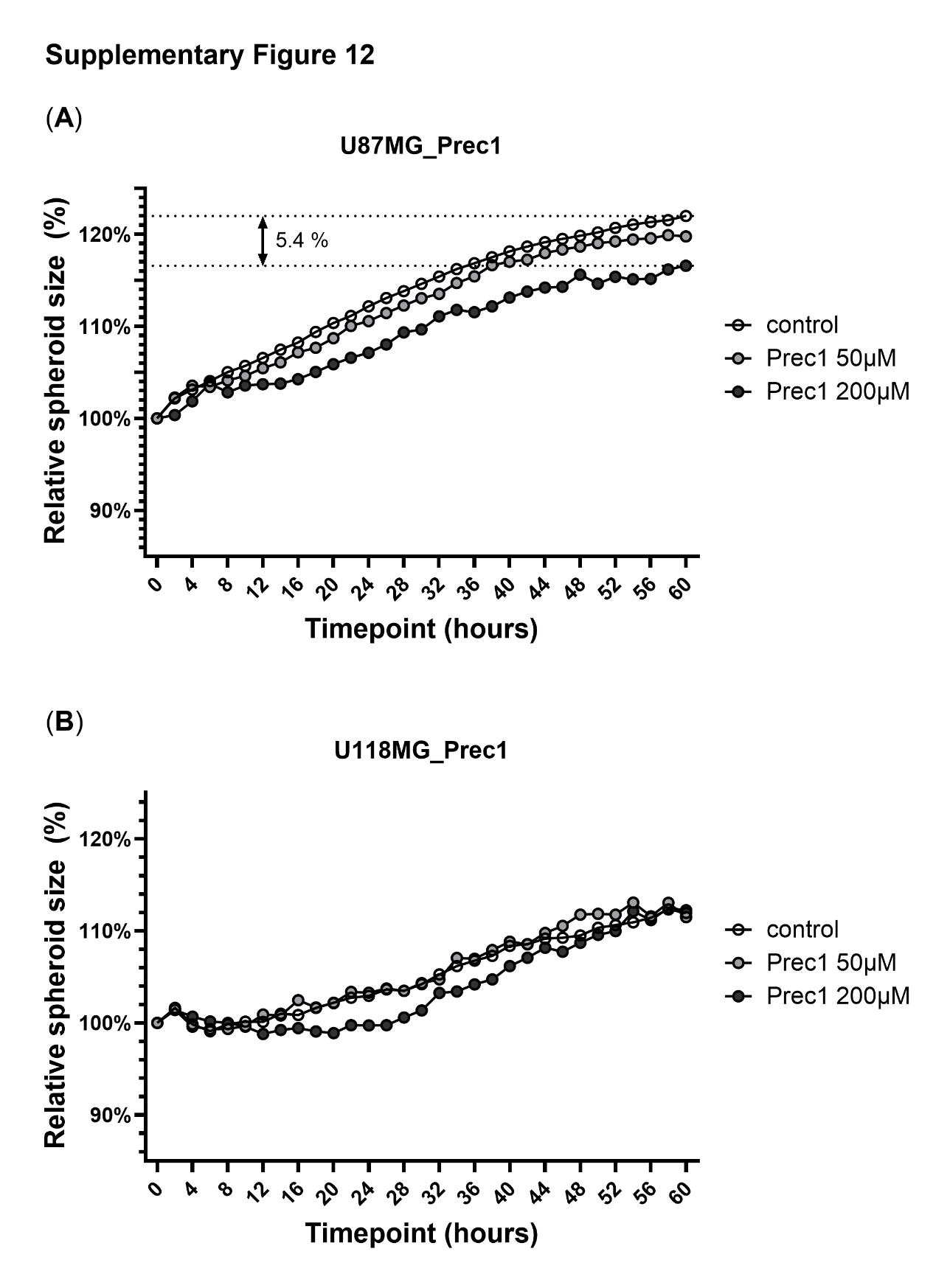

**Supplementary Figure 12.** Effect of conjugate precursor **Prec1** on U87MG (A) and U118MG (B) spheroid growth dynamics. n=2

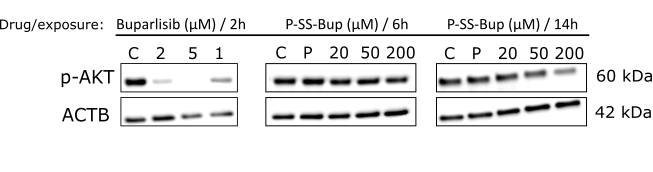
**Supplementary Figure 13.** Effect of Bup (2h exposure) and **P-SS-Bup**  (6 and 14h exposure) on the level of phosphorylated AKT protein (p-AKT) in U87MG cells under 2D conditions. Primary antibody p-AKT: p-Akt (Ser473); cat. no. #4060, Cell Signalling TECHNOLOGY^®^; loading control: β-actin (ACTB; cat. no. #A2228, Sigma-Aldrich). C = control (DMSO in CCM at a concentration corresponding to that 200 µM drug), P = Prec1 (200 µM).
